# Species-specific quorum sensing represses the chitobiose utilization locus in *Vibrio cholerae*

**DOI:** 10.1101/2020.04.16.045526

**Authors:** Catherine A. Klancher, Jane D. Newman, Alyssa S. Ball, Julia C. van Kessel, Ankur B. Dalia

## Abstract

The marine facultative pathogen *Vibrio cholerae* forms complex multicellular communities on the chitinous shells of crustacean zooplankton in its aquatic reservoir. *V. cholerae*-chitin interactions are critical for the growth, evolution, and waterborne transmission of cholera. This is due, in part, to chitin-induced changes in gene expression in this pathogen. Here, we sought to identify factors that influence chitin-induced expression of one locus, the chitobiose utilization operon (*chb*), which is required for the uptake and catabolism of the chitin disaccharide. Through a series of genetic screens, we identified that the master regulator of quorum sensing, HapR, is a direct repressor of the *chb* operon. We also found that the levels of HapR in *V. cholerae* are regulated by the ClpAP protease. Furthermore, we show that the canonical quorum sensing cascade in *V. cholerae* regulates *chb* expression in a HapR-dependent manner. Through this analysis we found that signaling via the species-specific autoinducer CAI-1, but not the inter-species autoinducer AI-2, influences *chb* expression. This phenomenon of species-specific regulation may enhance the fitness of this pathogen in its environmental niche.

**Importance:** In nature, bacteria live in multicellular and multispecies communities. Microbial species can sense the density and composition of their community through chemical cues using a process called quorum sensing (QS). The marine pathogen *Vibrio cholerae* is found in communities on the chitinous shells of crustaceans in its aquatic reservoir. *V. cholerae* interactions with chitin are critical for the survival, evolution, and waterborne transmission of this pathogen. Here, we show that *V. cholerae* uses QS to regulate the expression of one locus required for *V. cholerae*-chitin interactions.

## Introduction

The facultative bacterial pathogen *Vibrio cholerae*, the causative agent of the diarrheal disease cholera, natively resides in the aquatic environment. In this niche, *V. cholerae* forms multicellular communities on biotic and abiotic chitinous surfaces, like the shells of crustaceans or marine snow (1, 2). Chitin is a polysaccharide made up of β-1,4 linked *N*-acetylglucosamine (GlcNAc) and serves as a major nutrient source for *V. cholerae* in the marine environment (1, 3, 4). The ability of *V. cholerae* to form chitin biofilms is critical for the waterborne transmission of cholera (5, 6). As chitin is the most abundant biopolymer in the ocean, the ability of *Vibrio* species to break down and utilize this highly insoluble polysaccharide also serves an important role in global nitrogen and carbon recycling (1, 4).

When *V. cholerae* is associated with a chitinous surface, chitin induces the expression of a subset of genes in *V. cholerae*. The genes induced by chitin include those required for chitin degradation, uptake, and catabolism (termed the chitin utilization program), as well as the genes required for natural transformation (7, 8). Transcriptional responses resulting from *Vibrio*-chitin interactions are highly regulated. One major chitin-responsive regulator is the orphan hybrid sensor kinase ChiS (7, 9). ChiS senses chitin indirectly through the periplasmic <u>c</u>hitin <u>b</u>inding <u>p</u>rotein (CBP) (9, 10). In the absence of chitin, CBP represses ChiS through interactions with its periplasmic domain (9, 10). In the presence of chitin, the CBP-chitin complex stimulates ChiS activity (9, 10). Thus, in the presence of chitin, ChiS is active and can facilitate expression of the chitin utilization program. Alternatively, ChiS can be genetically activated in the absence of chitin by deleting *cbp* (10, 11).

In the marine environment, *V. cholerae* not only senses chitin to modulate gene expression, but also the presence of other bacteria through a process termed “quorum sensing” (QS) (12). This is a process by which bacteria indirectly sense other microbes in their community via small diffusible molecules called autoinducers (AIs). AIs are sensed by cognate sensor proteins. *V. cholerae* encodes four AI sensors, although the autoinducer molecules that serve as inducing cues are only known for two of them (13). AI sensing allows for cell-density specific gene expression programs, which regulate “group” or “individual” behaviors (14). *V. cholerae* senses both chitin and AIs to regulate natural transformation on chitinous surfaces (15). Though a link between chitin utilization and quorum sensing has previously been suggested, it has not been directly studied (16).

To investigate regulation of the chitin utilization program in *V. cholerae*, most studies employ the chitobiose utilization operon (*chb*) (10, 11, 17, 18). The *chb* operon encodes the genes required for uptake and catabolism of the chitin disaccharide chitobiose, and is highly induced in the presence of chitin oligosaccharides (7). Several mechanisms of *chb* regulation have already been identified. ChiS is the master regulator of the chitin utilization program in *V. cholerae*, and we have recently shown this protein is a direct transcriptional activator required for induction of the *chb* locus (9, 10). Previous work has shown that carbon catabolite repression (CCR) can also play a role in regulating chitin responsive phenotypes including ChiS-dependent induction of *chb* and natural transformation (18, 19). In addition, our group has previously found that the cell division licensing factor SlmA plays an essential role in activating *chb* expression (11). Tight regulation via these diverse signaling systems may act to ensure that the chitin utilization program is only expressed under conditions in which it will provide a competitive advantage.

Here, we sought to identify additional regulators of *chb*. Through a number of genetic screens and complementary molecular methods, we show that quorum sensing is an additional regulatory system that tunes expression of a chitin utilization locus in *V. cholerae*.

## Results

### ClpA is identified in an unbiased screen for activators of P_chb_

To identify additional genes required for activation of the *chb* locus, we conducted a transposon mutant screen. This was carried out in a strain containing a chromosomally-integrated P_*chb*_-*lacZ* transcriptional reporter. As shown previously, induction of P_*chb*_ is dependent on the activity of the master regulator ChiS (7, 10, 11). In the absence of chitin, ChiS activity is repressed by CBP. In the presence of chitin, CBP repression of ChiS is relieved, which allows for ChiS-dependent activation of P_*chb*_. In addition to being induced by chitin, ChiS can be activated genetically in the absence of chitin by deleting *cbp* (10, 11). As chitin oligomers are prohibitively expensive, a Δ*cbp* mutation was used to induce ChiS-dependent P_*chb*_-*lacZ* expression in our genetic screen exactly as previously described (11). So, the starting genotype for our screen was a strain containing P_*chb*_-*lacZ* and a Δ*cbp* mutation. This strain formed blue colonies on X-gal containing plates and we screened for white colonies to identify putative activators that contribute to P_*chb*_ induction.

Of approximately 60,000 transposon mutants visually screened for loss of P_*chb*_-*lacZ* expression, one gene identified was *clpA* (2 unique transposon insertions). Other hits identified in this screen are listed in **Table S1**. To study the effect of *clpA* on P_*chb*_ activity moving forward, we utilized a previously described chromosomally-integrated P_*chb*_-GFP reporter (10, 11). Using this reporter, we found that a Δ*clpA* mutation resulted in a ∼3-fold decrease in P_*chb*_ expression relative to the parent (**Fig. 1**). Importantly, complementation of this strain with an ectopic copy of *clpA in trans* restored P_*chb*_ expression to parent levels (**Fig. S1**). ClpA is a AAA+ unfoldase that recognizes protein substrates, unfolds them, and feeds them into the ClpP protease where they are degraded (20). If ClpA was exhibiting its effect on P_*chb*_ expression as a part of the ClpAP machine, we hypothesized that a Δ*clpP* mutation should phenocopy a Δ*clpA* mutation. Indeed, Δ*clpP* and Δ*clpP* Δ*clpA* strains phenocopied a Δ*clpA* mutant for P_*chb*_ expression (**Fig. 1**). These results suggest that loss of the ClpAP protease decreases P_*chb*_ expression. As ClpAP degrades proteins, we hypothesized that ClpAP may indirectly promote activation of P_*chb*_ by degrading a repressor of the *chb* locus.

**Figure 1.**
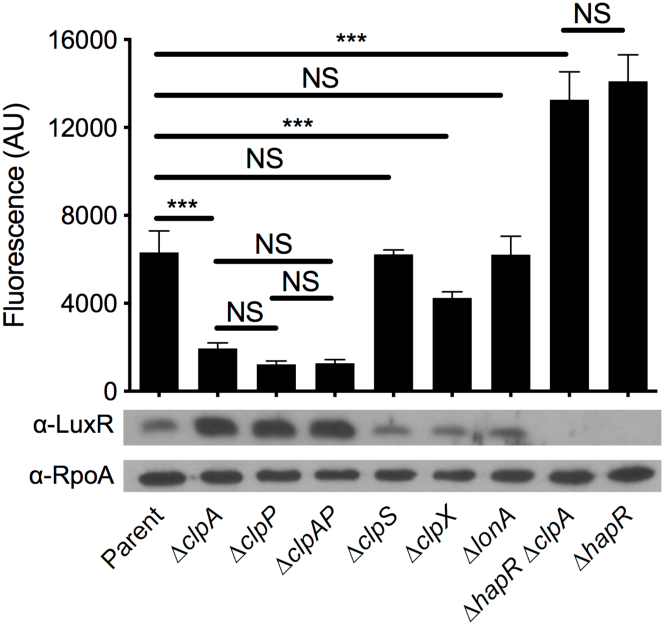
HapR is a repressor of P_chb_ that is degraded by the ClpAP protease. Expression of a P_*chb*_-GFP reporter and HapR protein levels were determined in the indicated mutant strains. The parent strain contained a P_*chb*_-GFP reporter and a Δ*cbp* mutation. A representative western blot is shown below bars to indicate the protein levels for HapR and RpoA (a loading control) in the corresponding strains. An antibody against LuxR, which has 72% identity and 86% similarity to HapR, is cross-reactive with HapR and so was used to detect HapR protein levels. Fluorescence of cultures was determined on a plate reader from at least six independent biological replicates and is shown as the mean ± SD. Statistical comparisons were made by one-way ANOVA with Tukey’s post-test. NS, not significant. ***, p < 0.001.

### HapR is a repressor of P_chb_ that is degraded by ClpAP

To identify a putative repressor of P_*chb*_ that is targeted by ClpAP for degradation, we conducted a counter-screen using the P_*chb*_-*lacZ* reporter. For the Δ*clpA* counter-screen, the parent strain had both Δ*cbp* and Δ*clpA* mutations. This mutant is white on X-gal plates because P_*chb*_-*lacZ* is poorly expressed; inactivation of the putative repressor in this strain should result in restoration of P_*chb*_-*lacZ* expression and yield a blue colony phenotype. In the Δ*clpA* counter-screen, we visually screened approximately 30,000 transposon mutants for re-activation of P_*chb*_-*lacZ* expression and identified *hapR* (9 unique transposon insertions). Other hits identified in this screen are listed in **Table S1**. A Δ*hapR* mutation restored P_*chb*_ expression in the Δ*clpA* mutant background (**Fig. 1**). In fact, a Δ*hapR* mutation allowed for higher P_*chb*_ expression than the parent strain. This suggests that HapR represses P_*chb*_ expression when ClpAP is intact and that ClpAP does not degrade the entire pool of HapR in the cell (**Fig. 1**). Importantly, the level of P_*chb*_ expression observed in the Δ*hapR* Δ*clpA* mutant phenocopied the Δ*hapR* strain (**Fig. 1**). This epistasis between *clpA* and *hapR* suggests that they are involved in the same pathway for regulating P_*chb*_ expression. In addition, complementation of the Δ*hapR* strain with an ectopic copy of *hapR in trans* decreased P_*chb*_ expression almost to the level in the parent (**Fig. S1**). Further, complementation of the Δ*hapR* Δ*clpA* strain with *hapR in trans* brought P_*chb*_ expression down to the level observed in the Δ*clpA* parent (**Fig. S1**).

We hypothesized that the reason P_*chb*_ expression was decreased in Δ*clpAP* mutants was due to increased HapR protein levels. Western blotting in these backgrounds revealed that HapR levels were, indeed, increased in strains containing mutations to *clpA* and/or *clpP* (**Fig. 1**). ClpAP dependent degradation of HapR is not unique to *V. cholerae*, but has previously been observed in *Vibrio vulnificus* where ClpAP degrades the HapR homolog SmcR (21). In addition to the ClpAP machine, it was shown that the Lon protease also plays a role in SmcR degradation. So, we next sought to investigate the role of other protease machines on induction of P_*chb*_ and HapR protein levels. Mutations to other Clp components (the ClpS adaptor protein or the ClpX unfoldase) did not have a marked impact on HapR protein levels. Also, a Δ*clpS* mutation did not affect P_*chb*_ expression levels, while a Δ*clpX* mutation slightly decreased P_*chb*_ expression (**Fig. 1**). Because HapR expression was not affected by the Δ*clpX* mutation, the observed decrease in P_*chb*_ expression may be attributed to a pleiotropic effect (*i.e.*, an effect of *clpX* that is independent of HapR-dependent P_*chb*_ repression). In contrast to the effect of the Lon protease on SmcR levels in *V. vulnificus*, we did not observe an impact of Δ*lonA* on HapR levels in *V. cholerae*; and Δ*lonA*, correspondingly, did not affect P_*chb*_ expression (**Fig. 1**). Together, these results establish that HapR is a repressor of P_*chb*_ and that HapR levels are controlled specifically by the ClpAP protease in *V. cholerae*.

### HapR-mediated repression of P_chb_ occurs on chitinous surfaces

Thus far, we have studied P_*chb*_ regulation using a Δ*cbp* mutation to induce ChiS activity. Natural induction of this locus, however, occurs in the presence of chitin oligosaccharides. So, we next wanted to test whether the repression of P_*chb*_ by HapR was observed in a more physiologically relevant setting. To test this, we cultured *V. cholerae* strains with CBP intact, a P_*chb*_-mCherry reporter, and a construct that constitutively expresses GFP (P_*const2*_-GFP) on chitin beads (**Fig. 2A**). The P_*const2*_-GFP construct (derived from the insulated *proD* promoter (22)) served as an internal control for noise in gene expression and was used to normalize P_*chb*_-mCherry expression in single cells (see Methods for details).

**Figure 2.**
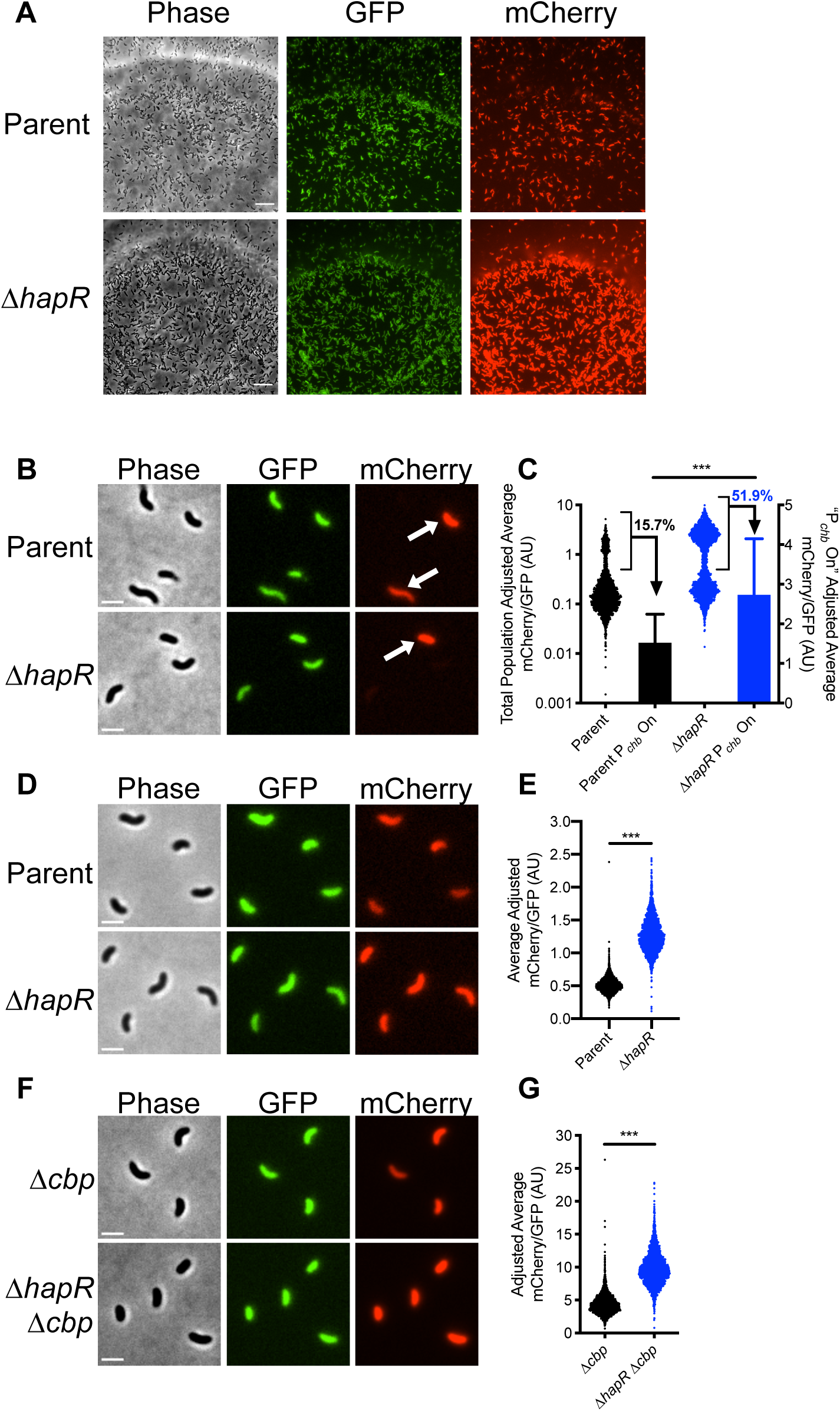
HapR-mediated repression of P_chb_ occurs on chitinous surfaces. **A**) Representative images of the indicated *V. cholerae* strains grown on chitin beads. The parent strain background contains both a P_*chb*_-mCherry reporter and a P_const2_-GFP construct, which exhibits constitutive GFP expression. Scale bar = 10 μm. **B-C**) P_*chb*_ expression in the indicated *V. cholerae cbp*^*+*^ strains grown on chitin beads. **B**) Representative images of cells that were vortexed off chitin beads. Arrows demarcate cells where P_*chb*_ expression is induced (“P_*chb*_ On” cells). Scale bar = 2 μm. **C**) Scatter plot and bar graph showing the relative expression of the P_*chb*_-mCherry reporter in cells cultured on chitin beads. Scatter plots (left Y-axis) represent the entire population, whereas the bars (right Y-axis) represent only the “P_*chb*_ On” cells (bracketed in black on scatter plots). The percentage of cells in the “P_*chb*_ On” population is indicated. n = 2120 for Parent; n = 2191 for Δ*hapR;* n = 333 for Parent P_*chb*_ On; n = 1139 for Δ*hapR* P_*chb*_ On. Data shown are from two independent biological replicates. **D-E**) P_*chb*_ expression of the indicated *V. cholerae* strains where P_*chb*_ expression is induced by chitin oligosaccharides. **D**) Representative images of cells grown with chitin oligosaccharides. Scale bar = 2 μm. **E**) Scatter plot showing the relative expression of a P_*chb*_-mCherry reporter in the indicated *V. cholerae* strains. n = 2735 for Parent; n = 2384 for Δ*hapR.* Data shown are from two independent biological replicates. **F-G**) P_*chb*_ expression of the indicated *V. cholerae* strains where P_*chb*_ expression is induced via deletion of *cbp*. **F**) Representative images of cells grown in the absence of chitin. Scale bar = 2 μm. **G**) Scatter plot showing the relative expression of a P_*chb*_-mCherry reporter in the indicated *V. cholerae* strains. n = 2407 for Δ*cbp*; n = 2313 for Δ*hapR Δcbp.* Data shown are from two independent biological replicates. Statistical comparisons in **D, G**, and **J** were made using Student’s t-test. ***, p < 0.001.

When cells were cultured on chitin beads, both the parent strain and the Δ*hapR* strain exhibited a bimodal distribution of P_*chb*_ expression (**Fig. 2A-C**). There were at least two possible explanations for the observed bimodality in P_*chb*_ gene expression in this experiment. One, the signaling pathway that leads to activation of *chb* has switch-like behavior that results in a population that exhibits bimodality in *chb* expression. Or two, the environment during growth on chitin beads is heterogeneous and only some cells within the population have access to the chitin oligosaccharides necessary for activation of *chb*. Those cells that do not have access to chitin do not activate P_*chb*_ expression (P_*chb*_ Off), while those that do have access to chitin do induce P_*chb*_ expression (P_*chb*_ On). To differentiate between these two possibilities, we induced expression of P_*chb*_ either using the native inducer, chitin oligosaccharides, or genetically by deleting *cbp*. If the signaling circuit responsible for P_*chb*_ activation has switch-like behavior, we would predict that bimodality in gene expression would be maintained in both of these conditions; however, if bimodality is the result of heterogeneous access to chitin oligosaccharides when cells are cultured on chitin beads, we would expect bimodality to be lost. When P_*chb*_ was induced by chitin oligosaccharides or *via* deletion of *cbp*, all cells in the population are uniformly “P_*chb*_ On” (**Fig. 2D, 2F**) and the population becomes unimodal (**Fig. 2E, 2G**). Thus, these results showed that only the “P_*chb*_ On” cells have access to chitin oligosaccharides when grown on chitin beads. These results are consistent with chitin-induction of natural transformation on chitin beads, where cells display bimodality in gene expression that is also likely due to heterogeneous access to inducing chitin oligosaccharides (23).

Next, we tested whether HapR influenced activation of P_*chb*_ on chitin beads by just assessing the expression level among “P_*chb*_ On” cells. As observed in bulk populations, the Δ*hapR* strain exhibited a ∼1.7 fold increase in P_*chb*_ expression when cultured on chitin beads (**Fig. 2C**) and a ∼2.5 fold increase when cultured with chitin oligosaccharides (**Fig. 2E**). These values were consistent with the ∼2-fold increase in P_*chb*_ expression observed in single cells when the population was induced via deletion of *cbp* (**Fig. 2G**). Finally, native *chb* transcripts (measured *via* qRT-PCR) were also induced ∼2-fold higher in a Δ*hapR* mutant compared to the parent when strains were induced with chitin oligosaccharides (**Fig. S2**). Together, these results indicate that HapR is a *bona fide* repressor of P_*chb*_ under physiologically relevant inducing conditions.

### HapR repression of P_chb_ is conserved in other V. cholerae El Tor isolates

Previously, it was suggested that HapR was an activator of P_*chb*_ expression in the *V. cholerae* El Tor isolate A1552 (16). Specifically, the authors showed that deletion of *hapR* resulted in a decrease in P_*chb*_ expression when cells were cultured on chitin flakes. In the present study, we use the *V. cholerae* El Tor isolate E7946 to study P_*chb*_ regulation. To assess if HapR exhibits different effects on P_*chb*_ expression depending on the strain background, we assessed expression of a P_*chb*_-mCherry reporter in *hapR*^+^ and Δ*hapR* derivatives of both E7946 and A1552. Consistent with our previous results, we observed that P_*chb*_ expression is elevated ∼2-fold in the Δ*hapR* derivative of both strain backgrounds when induced with chitin oligosaccharides or via deletion of *cbp*, which is consistent with HapR acting as a repressor of this locus (**Fig. S3**). It is unclear what explains the discrepancy between our findings and those that were previously reported; however, these data suggest that they cannot be attributed to differences between strain backgrounds.

### Deletion of HapR does not confer a growth advantage during growth on chitobiose

Our data indicates that in the absence of HapR, P_*chb*_ expression is elevated. The *chb* locus encodes the genes required for uptake and degradation of the chitin disaccharide, chitobiose. Thus, we wanted to assess if the increased expression of P_*chb*_ confers a fitness advantage to Δ*hapR* cells during growth on chitobiose. To test this, we conducted competitive growth assays with a 1:1 mixture of a parent and a Δ*hapR* mutant strain on minimal medium with chitobiose as the sole carbon source. We hypothesized that if the Δ*hapR* mutant had a competitive growth advantage due to increased expression of *chb*, then it should outcompete the parent strain in this assay. Even after ∼48 generations of growth on chitobiose, however, we did not observe a competitive advantage for the Δ*hapR* mutant (**Fig. S4**). HapR is a global regulator that controls the expression of dozens of genes (24). Thus, even though a Δ*hapR* mutant has increased P_*chb*_ expression, there may be negative pleiotropic effects associated with the Δ*hapR* mutation that masks any competitive advantage of derepression of P_*chb*_ in this mutant during growth on chitobiose.

### HapR binds P_chb_ in vitro and in vivo

Thus far, our data suggest that HapR is a repressor of P_*chb*_, but it does not distinguish whether HapR is a direct or indirect regulator of this locus. To assess if HapR could be regulating P_*chb*_ directly, we used an *in silico* approach to identify putative HapR binding sites in P_*chb*_ based on consensus binding sequences generated for the HapR homologs LuxR (from *Vibrio harveyi*) and SmcR (from *V. vulnificus*) via chromatin immunoprecipitation sequencing (ChIP-seq) (25, 26). Using the Motif Alignment and Search Tool (MAST) in the Multiple Em for Motif Elicitiation (MEME) Suite (27), we identified two potential HapR binding sites in P_*chb*_ (**Fig. 3A**). Interestingly, these binding sites (BSs) overlap with other elements in the *chb* promoter required for transcriptional activation. HapR BS 1 overlaps with the SlmA binding site, which is a critical activator of P_*chb*_ expression (11). HapR BS 2 overlaps with the −35 signal, which is required for RNA polymerase to bind to the promoter and initiate transcription. This sequence analysis suggests that the repressive effect of HapR may be due to HapR binding directly antagonizing SlmA and/or RNAP at this locus.

**Figure 3.**
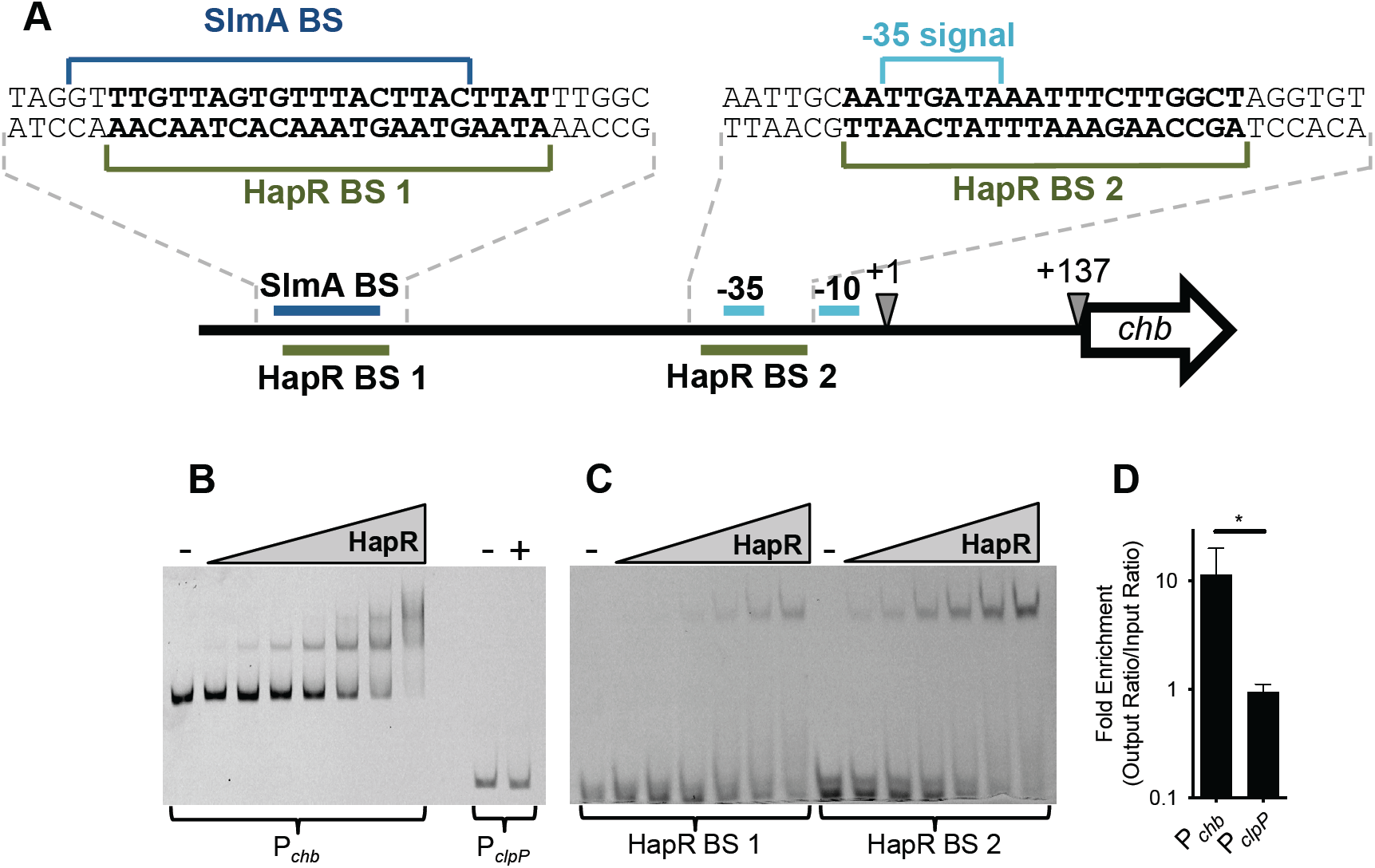
HapR binds P_chb_ in vitro and in vivo. **A**) Promoter map of P_*chb*_ highlighting putative HapR binding sites (BSs). Other sites required for P_*chb*_ activation (the SlmA BS and the −35 and −10 signals) are highlighted. The exact sequences of the region containing the HapR BSs (bolded), the SlmA BS, and the −35 signal are shown. The transcriptional start site (+1) and the translational start site (+137) are also shown. **B**) A representative EMSA using HapR and Cy5-labeled DNA probes of the indicated promoters. The P_*chb*_ probe was incubated with (from left to right) 0 nM (-), 12.5 nM, 25 nM, 50 nM, 100 nM, 200 nM, 400 nM, or 800 nM purified HapR. The P_*clpP*_ probe was incubated with 0 nM (-) or 800 nM HapR (+). **C**) A representative EMSA using HapR and 32 bp Cy5-labeled probes encompassing each of the putative HapR binding sites within the *chb* promoter (exact probe sequences are shown in **A**). The 32 bp probes were incubated with (from left to right) 0 nM (-), 100 nM, 200 nM, 400 nM, 800 nM, 1.6 µM, or 3.2 µM purified HapR. **D**) ChIP-qPCR assays showing enrichment of the indicated promoters relative to *rpoB*, a reference locus that HapR does not bind to. Data are from five independent biological replicates and shown as the mean ± SD. Statistical comparisons were made by Student’s t-test. *, p = 0.0240.

We next sought to determine whether HapR could directly bind to P_*chb*_ using both *in vitro* and *in vivo* approaches. First, we tested whether HapR could bind to P_*chb*_ *in vitro* using EMSA assays. Consistent with the presence of two HapR binding sites in P_*chb*_, we observed two shifts by EMSA when using a DNA probe of the *chb* promoter (**Fig. 3B**). HapR does not regulate the ClpP promoter and thus did not bind P_*clpP*_ (**Fig. 3B**), which is consistent with previous studies (28). EMSAs were also done using 32 bp probes that encompassed each putative HapR BS (probe sequences in **Fig. 3A**). We observed that HapR was able to bind to both probes, suggesting that HapR binds to both HapR BS 1 and HapR BS 2 (**Fig. 3C**). Next, we wanted to assess if HapR bound to P_*chb*_ *in vivo via* ChIP assays under physiologically relevant conditions. To that end, we first generated a FLAG-HapR strain that was functional for regulating P_*chb*_ expression (**Fig. S5**). Using this strain in ChIP-qPCR assays, we found that P_*chb*_ was, indeed, bound by HapR *in vivo*, while the negative control P_*clpP*_ locus was not bound by HapR (**Fig. 3D**). Together, these data demonstrate that HapR binds to P_*chb*_, which suggests that it is a direct repressor of this locus.

### Quorum sensing regulates expression of P_chb_ through the cholera-specific autoinducer CAI-1

HapR is the master regulator of quorum sensing (QS) in *V. cholerae*, and is highly expressed at high cell density (HCD) (29). Thus far, we have established that HapR is a repressor of P_*chb*_ expression. Next, we wanted to probe the role of QS in regulating P_*chb*_. To that end, we sought to test the effect of mutations in genes upstream of HapR in the *V. cholerae* QS cascade on P_*chb*_ induction and HapR protein levels.

QS in *Vibrio* species is controlled by autoinducer-responsive sensor proteins that indirectly modulate phosphorylation of the response regulator LuxO, which in turn indirectly regulates production of HapR (14). When autoinducer concentrations are low (*i.e.*, at low cell density (LCD)), multiple histidine kinase sensors acts as kinases (30-32). This results in high levels of phosphorylated LuxO, which prevents HapR production (**Fig. S6A**) (33). By contrast, at high autoinducer concentrations (*i.e.*, at HCD), the sensors act as phosphatases, which ultimately leads to dephosphorylation of LuxO and allows for HapR production (**Fig. S6B**) (14, 34).

First, we assessed the impact of HCD on P_*chb*_ induction and HapR levels by deleting *luxO*, which genetically locks cells in a HCD state. Induction of the parent strain is tested under HCD conditions, thus, as expected, HapR levels were similar between the parent and the Δ*luxO* mutant; accordingly, expression of P_*chb*_ was also similar between the parent and Δ*luxO* (**Fig. 4**). Next, we tested the effect of LCD on P_*chb*_ and HapR expression by generating a *luxO*^*D47E*^ mutant, which mimics phosphorylated LuxO and genetically locks cells in a LCD state. In this strain, we saw that P_*chb*_ expression increased and was correlated with a decrease in HapR protein levels (**Fig. 4**). Together, these data suggest that HapR repression of P_*chb*_ is mediated through the canonical QS circuit.

**Figure 4.**
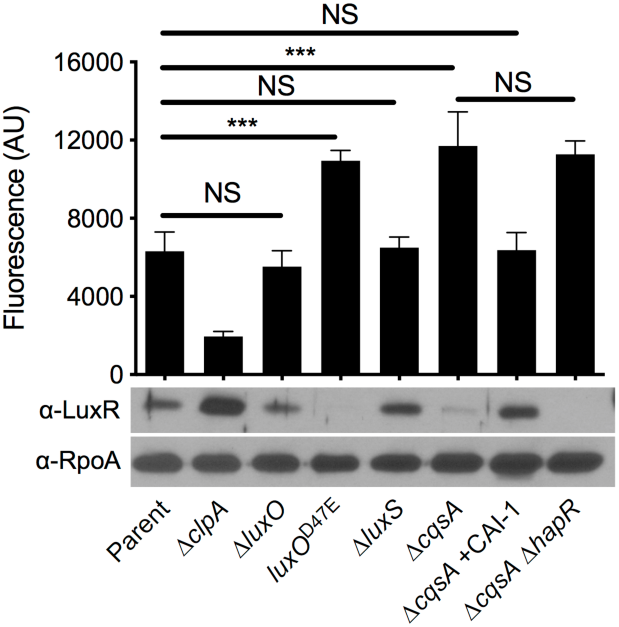
Quorum sensing regulates expression of P_chb_ through the cholera-specific autoinducer CAI-1. **A**) Model of QS regulation in *V. cholerae*. There are four sensor kinases that contribute to canonical QS in *V. cholerae* to regulate HapR production: LuxQ, CqsS, VpsS, and CqsR. The signals for VpsS and CqsR are unknown. LuxPQ senses the interspecies autoinducer AI-2 and CqsS senses the *V. cholerae*-specific autoinducer CAI-1. AI-2 is produced by the LuxS synthase and CAI-1 is produced by the CqsA synthase. (Left) At low cell density, the LuxQ and CqsS sensors act as kinases, resulting in phosphorylation of LuxO, which subsequently decreases expression of HapR. (Right) At high cell density when concentrations of AI-2 and CAI-1 are high, LuxQ and CqsS act as phosphatases, which results in dephosphorylation of LuxO, and subsequent activation of HapR expression. **B**) Expression of a P_*chb*_-GFP reporter and HapR protein levels were determined in the indicated strains. The parent strain contains a P_*chb*_-GFP reporter and a Δ*cbp* mutation. A representative western blot is shown below the bars to indicate the protein levels for HapR and RpoA (a loading control) in the corresponding strains. An antibody against LuxR, which has 72% identity and 86% similarity to HapR, is cross-reactive with HapR and so was used to detect HapR protein levels. Statistical comparisons were made by one-way ANOVA with Tukey’s post-test. NS, not significant. ***, p < 0.001. The P_*chb*_-GFP fluorescence data for “Parent” and “Δ*clpA*” are identical to the data presented in **Fig. 1** and were included here to ease comparisons.

Next, we wanted to move further upstream in the QS regulatory cascade to address if distinct autoinducers differentially affected expression of P_*chb*_ and HapR. There are four parallel histidine kinase sensors that coordinate QS-dependent control of HapR expression (13); however, the autoinducer signal is only known for two of these systems. The sensor LuxPQ is responsive to the inter-species autoinducer AI-2, and the sensor CqsS is responsive to the *V. cholerae*-specific autoinducer CAI-1 (**Fig. S6**) (35). To assess the role of each autoinducer in regulation of P_*chb*_, we made mutations to the synthase genes responsible for production of each autoinducer. LuxS makes AI-2 (36) and CqsA makes CAI-1 (35) (**Fig. S6**). In a strain that no longer produces AI-2 (Δ*luxS*), expression of P_*chb*_ was similar to that observed in the parent and HapR levels remained similar in these two strains (**Fig. 4**). By contrast, a strain that is unable to produce CAI-1 (Δ*cqsA*) had increased expression of P_*chb*_, likely due to the low level of HapR produced (**Fig. 4**). The observed decrease in P_*chb*_ expression in the Δ*cqsA* background was, indeed, due to a lack of CAI-1 production as exogenously adding back synthetic CAI-1 restored HapR levels and repression of P_*chb*_ to the parent level (**Fig. 4**). In addition, a strain that does not make CAI-1 induced P_*chb*_ to the same level as a strain that does not produce both CAI-1 or HapR (**Fig. 4**). This epistasis indicates that CAI-1 production and HapR are involved in the same regulatory pathway for modulating expression of P_*chb*_. These data support previous results, which indicate that CqsS signaling plays a dominant role in regulating HapR (23, 35, 37, 38). Together, these results indicate that P_*chb*_ expression is strongly influenced by QS signaling mediated by the *V. cholerae*-specific autoinducer CAI-1 and less via the inter-species autoinducer AI-2.

## Discussion

Here, we show that HapR acts as a repressor of chitobiose utilization genes in *V. cholerae*. When this organism forms communities on a chitinous surface, chitin induces expression of P_*chb*_ through activation of the chitin sensor ChiS (**Fig. 5**). *V. cholerae* then modulates P_*chb*_ expression depending on the presence of other *V. cholerae* cells, which it senses via the *V. cholerae*-specific autoinducer CAI-1. At low CAI-1 concentrations, P_*chb*_ expression is high because the repressor HapR is produced at a low level (**Fig. 5A**). At high CAI-1 concentrations, HapR is produced at higher levels and can repress P_*chb*_ expression (**Fig. 5B**). As the HapR binding sites in P_*chb*_ overlap with the SlmA binding site and the −35 signal, it is possible that HapR competes with these activators for binding at P_*chb*_. Thus, HapR-mediated repression may occur through antagonism of the binding activity of SlmA and/or RNAP at P_*chb*_.

**Figure 5.**
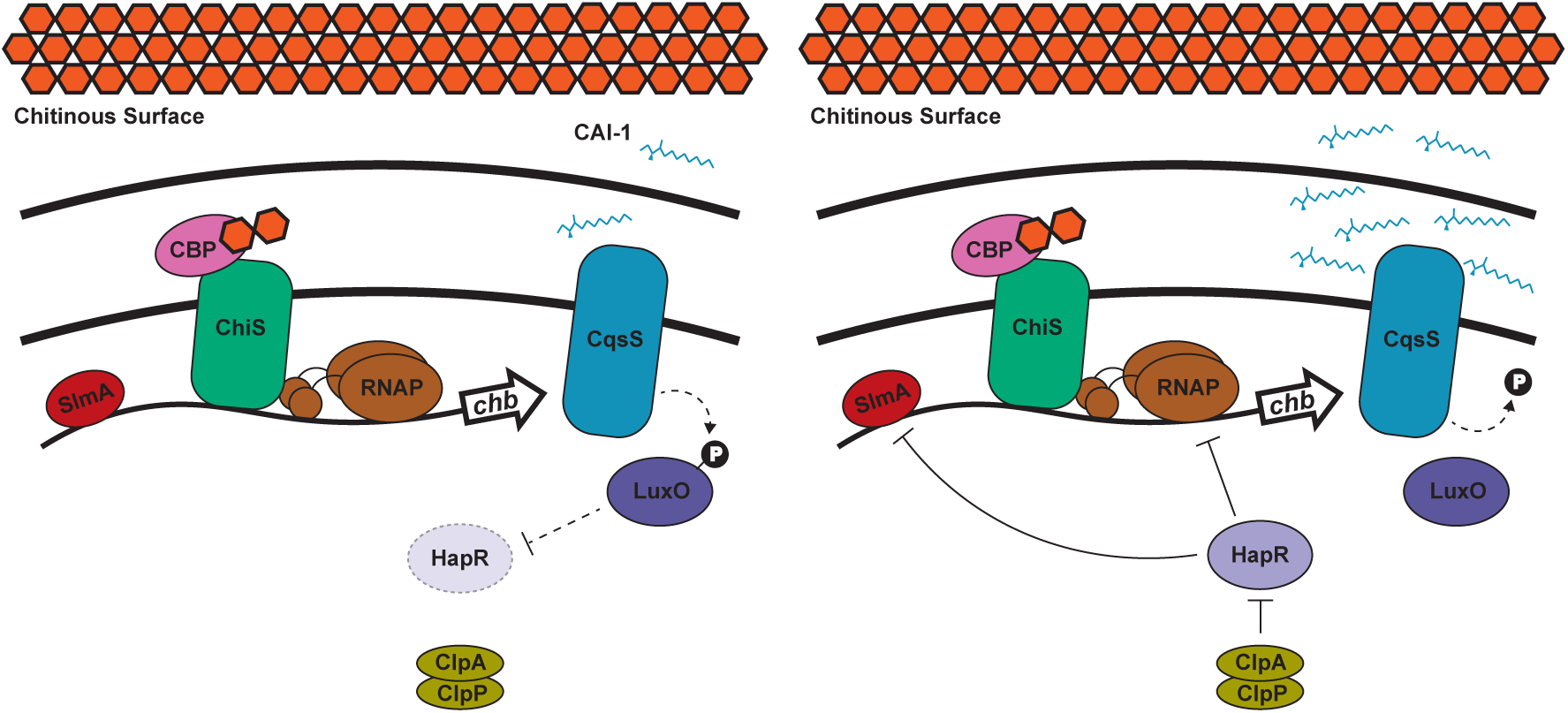
Model of quorum sensing regulation of chitobiose utilization genes in V. cholerae. When *V. cholerae* grows on a chitinous surface, the periplasmic chitin binding protein (CBP) binds to chitin oligosaccharides. This de-represses ChiS, which can then activate expression of the chitobiose utilization operon (*chb*) through recruitment of RNA Polymerase (RNAP) (10). Binding of SlmA to P_*chb*_ is also required for transcriptional activation (11). **A**) At low CAI-1 concentrations, the CqsS sensor indirectly (dashed line) promotes LuxO phosphorylation, which indirectly blocks HapR expression. In this state, P_*chb*_ is maximally expressed. **B**) At high CAI-1 concentrations, the CqsS sensor indirectly dephosphorylates LuxO, which results in indirect activation of HapR expression. HapR then exerts a repressive effect on P_*chb*_ by decreasing expression of this locus ∼2-fold in the presence of the ClpAP protease machine. If HapR is not degraded by ClpAP, its repressive effect can result in a ∼7-fold decrease in P_*chb*_ expression. The mechanism of HapR repression may be through occluding binding of SlmA or RNAP to P_*chb*_.

When HapR is produced, it gets proteolyzed by the ClpAP machine (**Fig. 5B**). While the dynamic range of HapR-dependent P_*chb*_ repression is only ∼2-fold when ClpAP is intact, the dynamic range of repression increases to ∼7-fold in the absence of ClpAP. Thus, it is tempting to speculate that regulation of ClpAP may modulate P_*chb*_ expression under some conditions. Currently, however, little is known about the regulation of ClpAP in *V. cholerae*. ClpAP is induced during heat shock in *V. vulnificus* (21). Also, in *E. coli*, ClpP expression is induced during heat shock (39-41). Thus, it is believed that the ClpP protease plays a role in the heat shock response. In *Bacillus subtilis*, ClpP is upregulated under various stress conditions, suggesting that this protease may also be induced by a general stress response (42). Thus, one possibility is that stress-dependent regulation of ClpAP indirectly regulates P_*chb*_. It has also been hypothesized that *V. cholerae* makes use of proteolysis machines to rapidly respond to changes in their environment; namely, the transition from the human gut to the aquatic environment after infection (43). ClpAP acting as an activator of P_*chb*_ may be a way for cells to rapidly induce expression of a locus that is not important in the human gut, but is useful for survival in its aquatic reservoir. Consistent with this idea, *chb* is induced late in infection, suggesting this is a critical locus for preparing to re-enter the aquatic environment after infection of a host (44).

The mechanisms underlying distinct responses to AI-2 or CAI-1 remain a topic of interest in QS. It has been shown that the CAI-1 sensor CqsS plays a dominant role over LuxPQ in modulating HapR levels. Thus, it remains possible that the presence of CAI-1 allows for more robust regulation of HapR-regulated genes. CAI-1 has been shown to be critical for expression of the virulence factor *hapA* (38), natural transformation (15), and for repressing chitobiose utilization as shown in this study. HapA is a protease that has been implicated in mediating *V. cholerae* detachment from host epithelial cells, thereby promoting dissemination of cells back into the aquatic environment (45, 46); natural transformation and chitin utilization aid in *V. cholerae* fitness in the marine environment. By contrast, expression of *tcpA*, a protein which contributes to *V. cholerae* pathogenesis in the human gut, was shown to be primarily regulated by AI-2 (37). It is tempting to speculate that CAI-1 controls *V. cholerae* behaviors important for marine survival, whereas AI-2 controls behaviors involved in infection. It has been hypothesized that sensing both of these autoinducers plays a critical role in biofilm dispersal, and that only when both are sensed, *V. cholerae* cells are induced to leave a surface (*i.e.* QS works as a coincidence detector for both signals) (37). Thus, signaling *via* distinct autoinducers may allow *V. cholerae* to modulate responses depending on the context of the environment they inhabit.

The data we present here suggests that at HCD, *V. cholerae* dampens its expression of a chitin utilization locus. Below, we speculate on a few reasons why this regulation may be advantageous. Chitin polymers in the shells of crustacean zooplankton are long chain polysaccharides in a crystalline insoluble lattice. In order to be used as a nutrient source, the long chain chitin must be broken down into smaller, soluble chitin oligosaccharides. *V. cholerae* secretes chitinases that enzymatically degrade insoluble chitin into soluble chitin oligosaccharides for uptake and catabolism. The production and secretion of chitinases is an energetically costly process; thus, liberated chitin is a valuable “public good” in the context of a chitin biofilm (47). So, it is possible that QS regulation of chitin utilization allows *V. cholerae* to modulate chitin uptake based on the composition of its microbial community. When the level of *V. cholerae* cells (and the corresponding concentration of cholera specific autoinducer CAI-1) in the community is high, chitin uptake may decrease among individual cells within the population in an effort to “share” liberated chitin oligosaccharides. By contrast, when the level of *V. cholerae* in the community is low, sensing of only the inter-species signal AI-2 (which is produced by many bacterial species) does not suppress chitin uptake and utilization, thus, allowing the *V. cholerae* within this population to maximally compete for liberated chitin oligosaccharides.

Another possibility is that this regulation allows *V. cholerae* to control its production of toxic metabolites. In a previous study, it was shown that cells in a HCD state alter their metabolic flux to produce neutral byproducts as opposed to organic acids when grown in LB supplemented with a fermentable sugar like glucose (48). This regulation allows for a more stable community; by contrast, cells locked in a LCD state will excrete harmful metabolic byproducts, which leads to the demise of the community (48). It has been shown that *V. cholerae* excretes ammonium as a potentially toxic byproduct when grown on chitin (16). Also, because chitin oligosaccharides likely feed into glycolysis, it is possible that catabolism of chitin results in the production of potentially toxic organic acids.

Thus, repressing chitin uptake and utilization may also help slow the rate of metabolism to prevent the accumulation of toxic intermediates in a dense community setting.

## Materials & Methods

### Bacterial strains and culture conditions

*V. cholerae* strains were routinely grown at 30°C in LB medium and on LB agar supplemented when necessary with carbenicillin (20 μg/mL or 50 μg/mL), kanamycin (50 μg/mL), spectinomycin (200 μg/mL), trimethoprim (10 μg/mL), and/or chloramphenicol (2 μg/mL). Strains were grown in DASW medium for microscopy (see below for details), Instant Ocean for generating mutant strains (see below for details), or M9 minimal medium for competition experiments (see below for details).

### Transposon mutagenesis

Transposon mutant libraries were generated with a Carb^R^ mini-Tn10 transposon exactly as previously described (49). Briefly, the transposon mutagenesis plasmid pDL1086 was first mated into parent strains containing chromosomally-integrated P_*chb*_-*lacZ* and a Δ*cbp* mutation (activator screen) or Δ*clpA* Δ*cbp* mutations (Δ*clpA* counter-screen). The parent strains also carried a chromosomally-integrated P_*chb*_-mCherry reporter at an ectopic site to ensure that candidate transposon mutants affected expression of P_*chb*_ and did not simply disrupt the P_*chb*_-*lacZ* reporter. The activator screen parent also carried an additional copy of ChiS at an ectopic site, which prevented transposon hits in this known regulator of P_*chb*_. Transposition was induced by plating cultures on LB agar supplemented with 50 μg/mL Carb at 42°C. To screen colonies, plates also contained 40 μg/mL X-gal and 5 mM IPTG. IPTG was added to competitively inhibit the basal activity of the P_*chb*_-*lacZ* reporter.

The sequence of transposon-genomic junctions in transposon mutants were determined by inverse PCR followed by Sanger sequencing. Briefly, genomic DNA was purified from mutants of interest and digested with the FatI restriction enzyme per manufacturer’s instructions (NEB). Digested genomic DNAs were then incubated with T4 DNA ligase per manufacturer’s instructions (NEB) to generate self-ligated circles. The transposon-genomic junction was then amplified by PCR using the primers specified in **Table 2** and subsequently Sanger sequenced (Eurofins Genomics).

**Table 1.** Strains used in this study.

| Strain Designation | Reference in Manuscript | Genotype | Reference |
| --- | --- | --- | --- |
| SAD 030 | *V. cholerae* E7946 WT; parent for all other E7946 strains; Fig. S2 Parent | *V. cholerae* E7946 O1 El Tor | (50) |
| SAD 2825 | Activator screen strain | *V. cholerae* E7946 ΔVC1807::P<sub>chb</sub>-mCherry, Kan<sup>R</sup>; P<sub>chb</sub>-lacZ, Spec<sup>R</sup>; ΔVCA0692::chiS, Tm<sup>R</sup>; Δ3' cbp, pDL1086 Cm<sup>R</sup> Carb<sup>R</sup> | This study |
| SAD 2826 | ΔclpA counter-screen strain | *V. cholerae* E7946 ΔVC1807::P<sub>chb</sub>-mCherry, Kan<sup>R</sup>; P<sub>chb</sub>-lacZ, Spec<sup>R</sup>; ΔclpA; Δcbp::Tm<sup>R</sup>; pDL1086, Cm<sup>R</sup> Carb<sup>R</sup> | This study |
| SAD 1309 | Fig. 1 & 4 Parent; Fig. S5 HapR WT | *V. cholerae* E7946 ΔlacZ::P<sub>chb</sub>-GFP, Kan<sup>R</sup>; Δcbp::Spec<sup>R</sup> | (11) |
| SAD 2827 | Fig. 1 & 4 ΔclpA | *V. cholerae* E7946 ΔlacZ::P<sub>chb</sub>-GFP, Kan<sup>R</sup>; ΔclpA::Carb<sup>R</sup>; Δcbp::Spec<sup>R</sup> | This study |
| SAD 2828 | Fig. 1 ΔclpP | *V. cholerae* E7946 ΔlacZ::P<sub>chb</sub>-GFP, Kan<sup>R</sup>; ΔclpP::Carb<sup>R</sup>; Δcbp::Tm<sup>R</sup> | This study |
| SAD 2829 | Fig. 1 ΔclpAP | *V. cholerae* E7946 ΔlacZ::P<sub>chb</sub>-GFP, Kan<sup>R</sup>; ΔclpA::Carb<sup>R</sup>; ΔclpP::Tm<sup>R</sup>; Δcbp::Spec<sup>R</sup> | This study |
| SAD 2830 | Fig. 1 ΔclpS | *V. cholerae* E7946 ΔlacZ::P<sub>chb</sub>-GFP, Kan<sup>R</sup>; ΔclpS::Carb<sup>R</sup>; Δcbp::Tm<sup>R</sup> | This study |
| SAD 2831 | Fig. 1 ΔclpX | *V. cholerae* E7946 ΔlacZ::P<sub>chb</sub>-GFP, Kan<sup>R</sup>; ΔclpX::Carb<sup>R</sup>; Δcbp::Tm<sup>R</sup> | This study |
| SAD 2832 | Fig. 1 ΔlonA | *V. cholerae* E7946 ΔlacZ::P<sub>chb</sub>-GFP, Kan<sup>R</sup>; ΔlonA::Tm<sup>R</sup>; Δcbp::Carb<sup>R</sup> | This study |
| SAD 2833 | Fig. 1 ΔhapR ΔclpA | *V. cholerae* E7946 ΔlacZ::P<sub>chb</sub>-GFP, Kan<sup>R</sup>; ΔhapR::Spec<sup>R</sup>; ΔclpA::Carb<sup>R</sup>; Δcbp::Tm<sup>R</sup> | This study |
| SAD 2834 | Fig. 1 ΔhapR | *V. cholerae* E7946 ΔlacZ::P<sub>chb</sub>-GFP, Kan<sup>R</sup>; ΔhapR::Spec<sup>R</sup>; Δcbp::Carb<sup>R</sup> | This study |
| SAD 2835 | Fig. 2 Parent | *V. cholerae* E7946 ΔlacZ::P<sub>chb</sub>-mCherry, Kan<sup>R</sup>; ΔVCA0692::P<sub>const2</sub>-GFP, Spec<sup>R</sup> | This study |
| SAD 2836 | Fig. 2 ΔhapR | *V. cholerae* E7946 ΔlacZ::P<sub>chb</sub>-mCherry, Kan<sup>R</sup>; ΔVCA0692::P<sub>const2</sub>-GFP, Spec<sup>R</sup>; ΔhapR::Cm<sup>R</sup> | This study |
| SAD 2837 | Fig. 2 Δcbp & Fig. S3 E7946 Parent | *V. cholerae* E7946 ΔlacZ::P<sub>chb</sub>-mCherry, Kan<sup>R</sup>; ΔVCA0692::P<sub>const2</sub>-GFP, Spec<sup>R</sup>; Δcbp::Tm<sup>R</sup> | This study |
| SAD 2838 | Fig. 2 ΔhapR Δcbp & Fig. S3 E7946 ΔhapR | *V. cholerae* E7946 ΔlacZ::P<sub>chb</sub>-mCherry, Kan<sup>R</sup>; ΔVCA0692::P<sub>const2</sub>-GFP, Spec<sup>R</sup>; ΔhapR::Cm<sup>R</sup>; Δcbp::Tm<sup>R</sup> | This study |
| SAD 2839 | Fig. 3 ChIP strain and Fig. S5 1x FLAG-HapR | *V. cholerae* E7946 ΔlacZ::P<sub>chb</sub>-GFP, Kan<sup>R</sup>; 1x FLAG-hapR; Δcbp::Tm<sup>R</sup> | This study |
| SAD 2840 | Fig. 4 ΔluxO | *V. cholerae* E7946 ΔlacZ::P<sub>chb</sub>-GFP, Kan<sup>R</sup>; ΔluxO::Spec<sup>R</sup>; Δcbp::Carb<sup>R</sup> | This study |
| SAD 2841 | Fig. 4 luxO<sup>D47E</sup> | *V. cholerae* E7946 ΔlacZ::P<sub>chb</sub>-GFP, Kan<sup>R</sup>; luxO<sup>D47E</sup>; Δcbp::Spec<sup>R</sup> | This study |
| SAD 2842 | Fig. 4 ΔluxS | *V. cholerae* E7946 ΔlacZ::P<sub>chb</sub>-GFP, Kan<sup>R</sup>; ΔluxS::Cm<sup>R</sup>; Δcbp::Spec<sup>R</sup> | This study |
| SAD 2843 | Fig. 4 ΔcqsA | *V. cholerae* E7946 ΔlacZ::P<sub>chb</sub>-GFP, Kan<sup>R</sup>; ΔcqsA::Tm<sup>R</sup>; Δcbp::Spec<sup>R</sup> | This study |
| SAD 2844 | Fig. 4 ΔcqsA ΔhapR | *V. cholerae* E7946 ΔlacZ::P<sub>chb</sub>-GFP, Kan<sup>R</sup>; ΔcqsA::Tm<sup>R</sup>; ΔhapR::Cm<sup>R</sup>; Δcbp::Spec<sup>R</sup> | This study |
| SAD 306 | *V. cholerae* A1552 WT; parent for all other A1552 strains | *V. cholerae* A1552 O1 El Tor | (51) |
| SAD 2908 | *E. coli* strain used to mate pMMB tfoX-qstR into complementation strains used in Fig. S1 | *E. coli* S17 harboring pMMB67EH tfoX qstR, Cm<sup>R</sup> | This study |
| SAD 2909 | Fig. S1 $\Delta hapR$ P<sub>hapR</sub>-hapR | *V. cholerae* E7946 $\Delta lacZ::P_{chb}$-GFP Kan<sup>R</sup>; $\Delta hapR::Spec^R$; $\Delta cbp::Carb^R$; $\Delta VCA0692::P_{hapR}$-hapR, Tm<sup>R</sup> | This study |
| SAD 2910 | Fig. S1 $\Delta hapR$ $\Delta clpA$ P<sub>hapR</sub>-hapR | *V. cholerae* E7946 $\Delta lacZ::P_{chb}$-GFP Kan<sup>R</sup>; $\Delta hapR::Spec^R$; $\Delta clpA::Carb^R$; $\Delta cbp::Tm^R$; $\Delta VCA0692::P_{hapR}$-hapR, Zeo<sup>R</sup> | This study |
| SAD 2911 | Fig. S1 $\Delta clpA$ P<sub>clpSA</sub>-clpSA | *V. cholerae* E7946 $\Delta lacZ::P_{chb}$-GFP Kan<sup>R</sup>; $\Delta clpA::Carb^R$; $\Delta cbp::Spec^R$; $\Delta VCA0692::P_{clpSA}$-clpSA, Tm<sup>R</sup> | This study |
| SAD 2912 | Fig. S2 $\Delta hapR$ | *V. cholerae* E7946 $\Delta hapR::Cm^R$ | This study |
| SAD 2845 | Fig. S3 A1552 Parent | *V. cholerae* A1552 $\Delta VCA0692::P_{const2}$-GFP, Spec<sup>R</sup>; $\Delta lacZ::P_{chb}$-mCherry, Kan<sup>R</sup>; $\Delta cbp::Tm^R$ | This study |
| SAD 2846 | Fig. S3 A1552 $\Delta hapR$ | *V. cholerae* A1552 $\Delta VCA0692::P_{const2}$-GFP, Spec<sup>R</sup>; $\Delta lacZ::P_{chb}$-mCherry, Kan<sup>R</sup>; $\Delta hapR::Cm^R$; $\Delta cbp::Tm^R$ | This study |
| SAD 2847 | Fig. S4 WT $lacZ^+$ | *V. cholerae* E7946 $\Delta VC1807::Cm^R$; $\Delta VC0995$ | This study |
| SAD 2848 | Fig. S4 WT $\Delta lacZ$ | *V. cholerae* E7946 $\Delta lacZ::Spec^R$; $\Delta VC1807::Cm^R$; $\Delta VC0995$ | This study |
| SAD 2849 | Fig. S4 $\Delta hapR$ $\Delta lacZ$ | *V. cholerae* E7946 $\Delta lacZ::Spec^R$; $\Delta VC1807::Cm^R$; $\Delta VC0995$; $\Delta hapR::Tm^R$ | This study |
| JDN 71 | Fig. 3B, 3C purified HapR | *E. coli* BL21(DE3) harboring pET28b-6XHis-hapR, Kan<sup>R</sup> | This study |

**Table 2.**
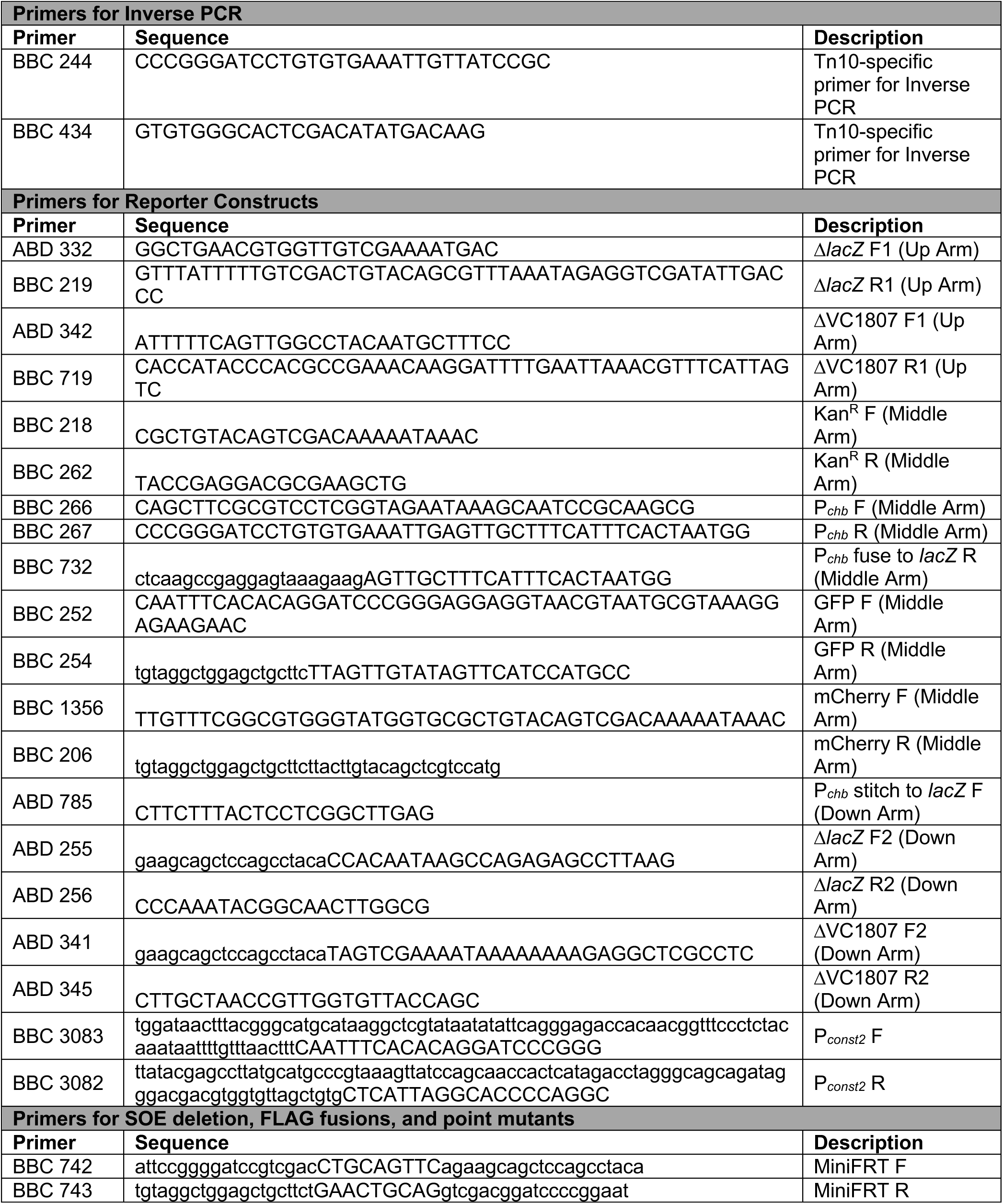
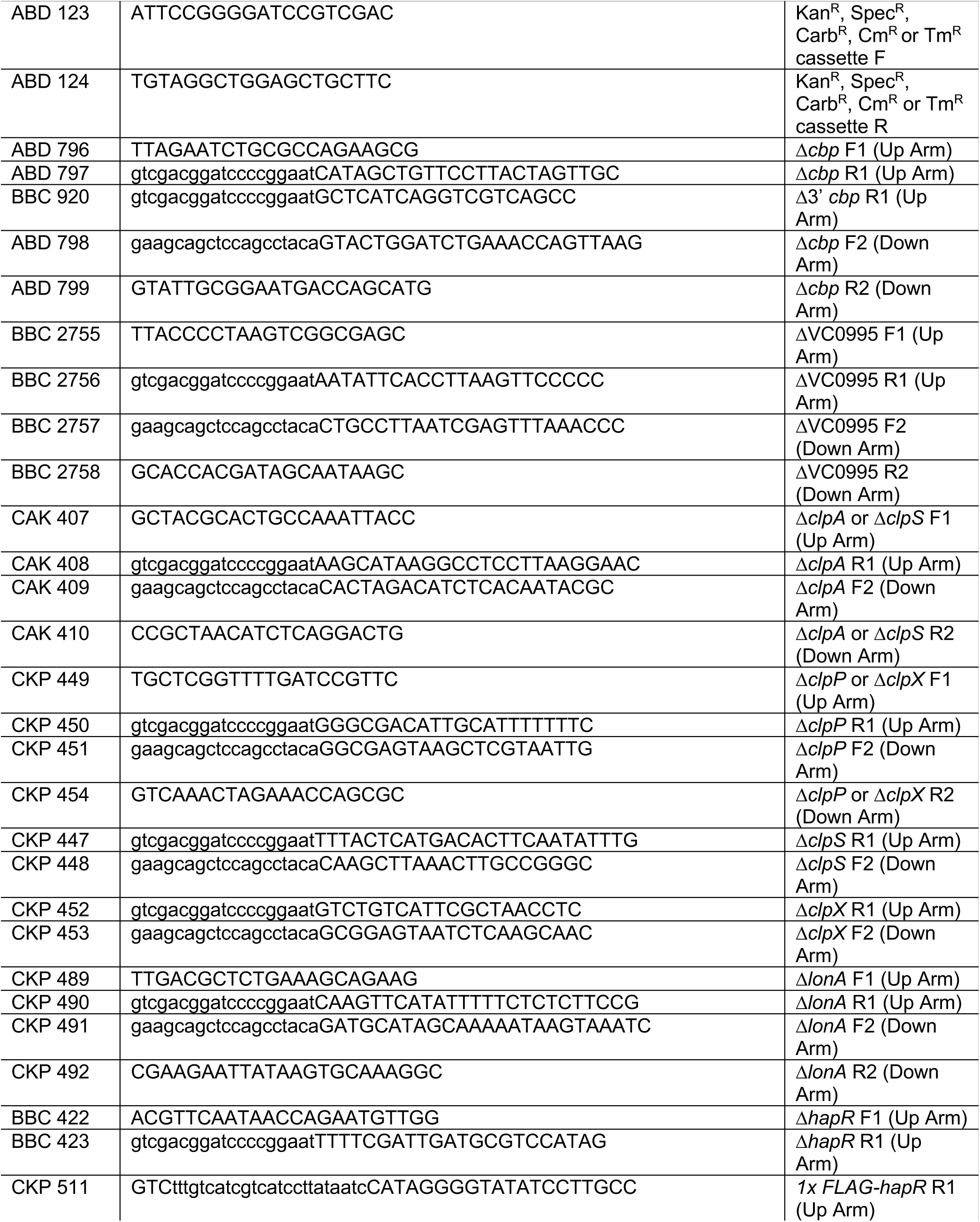
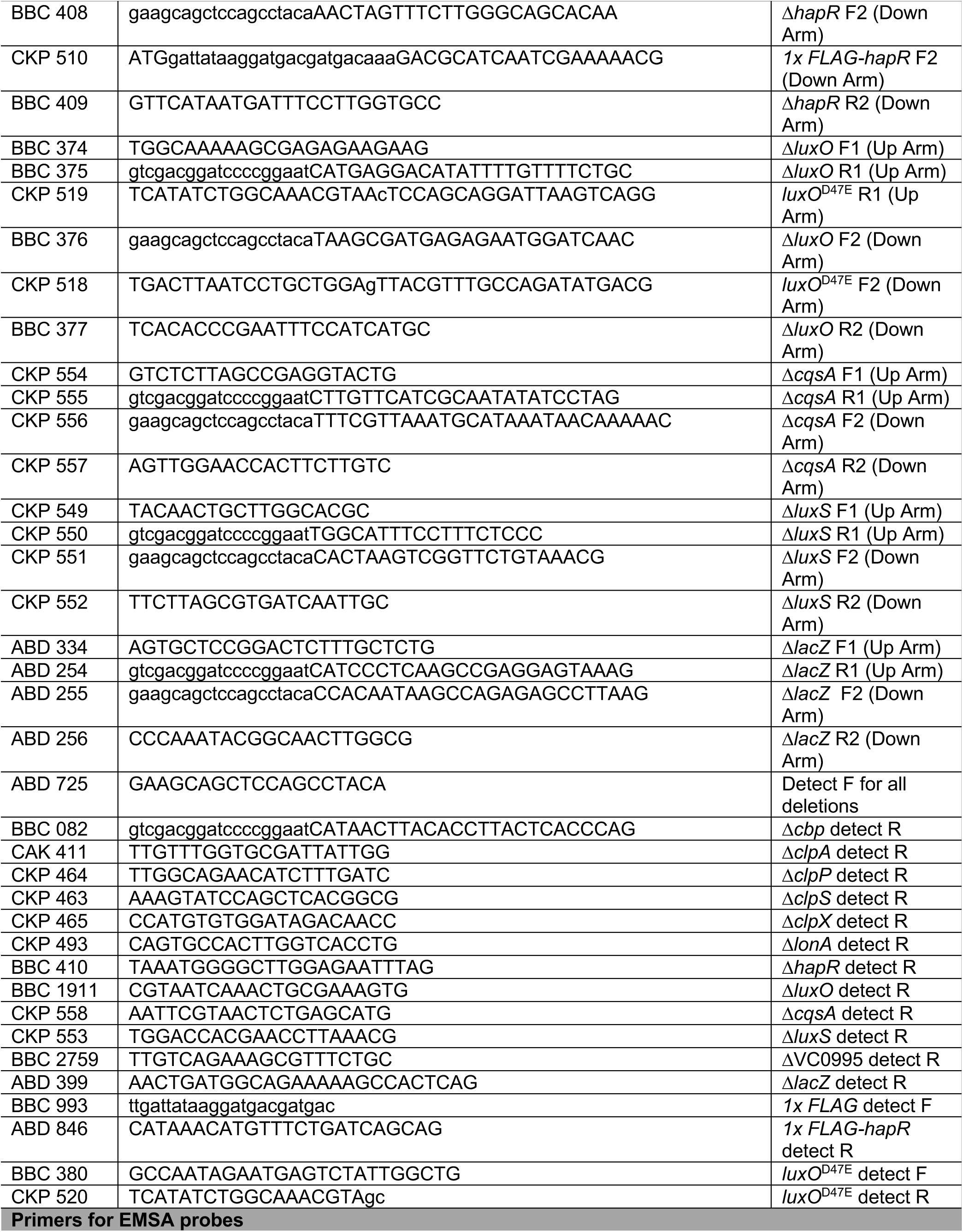
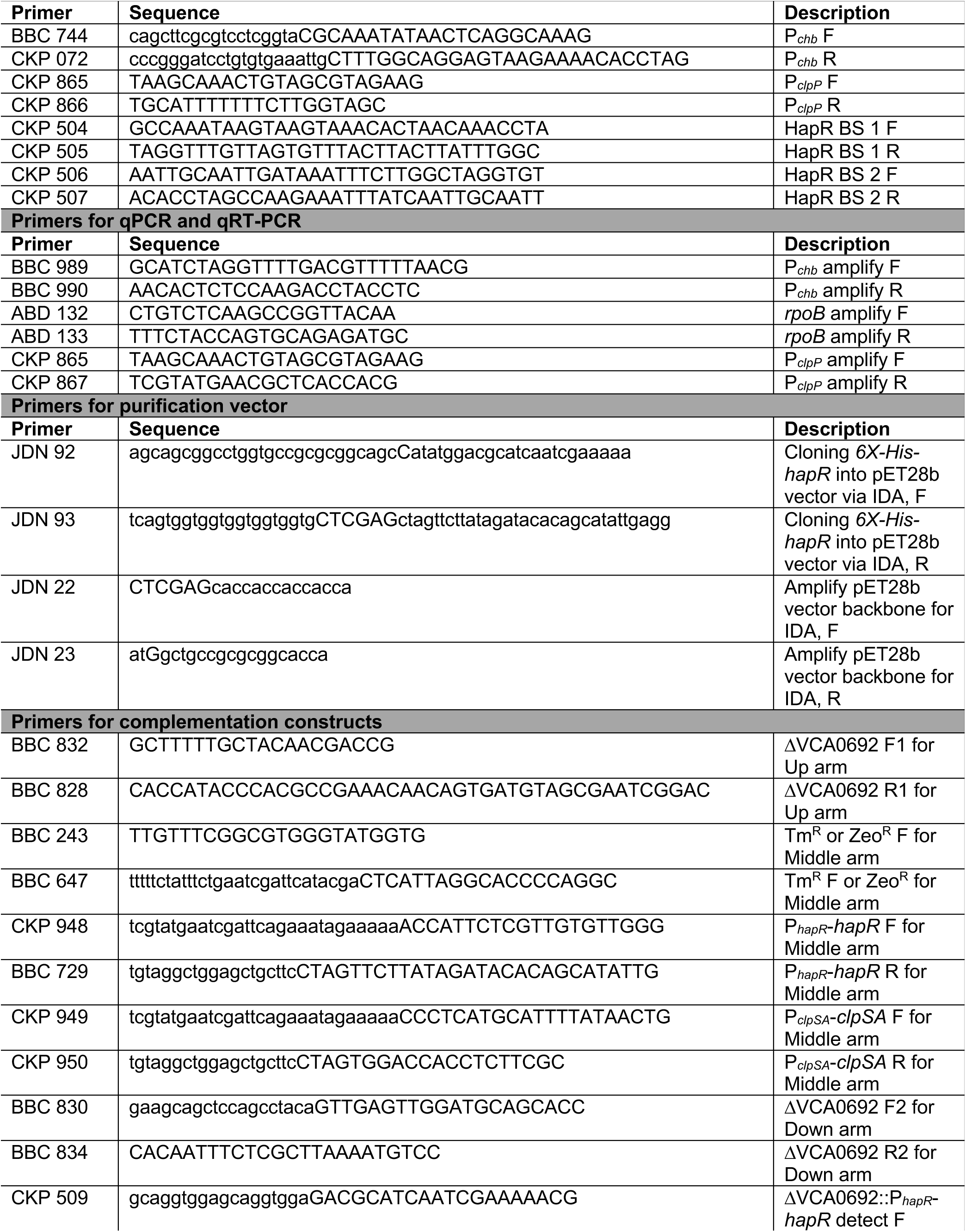
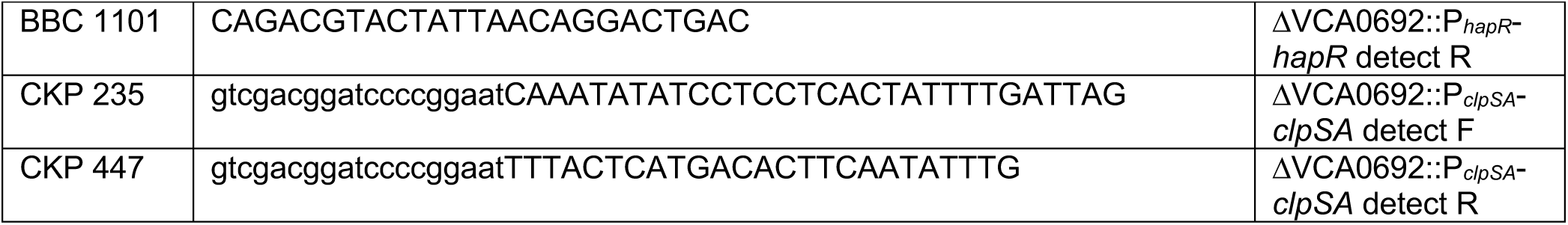
Primers used in this study.

### Generating mutant strains

*V. cholerae* E7946 served as the parent for all strains generated in this study, except for those used in **Fig. S3** where we compare E7946 to A1552 (50, 51). Mutant constructs were generated by splicing-by-overlap extension (SOE) PCR exactly as previously described with the primers indicated in **Table 2**. (52). PCRs were performed to generate Up (F1/R1), Middle, and Down (F2/R2) arms. Up and Down arms were designed to have 3kb arms of homology to the genome at the site targeted for mutagenesis. All three arms were then mixed and used as template for SOE PCR reactions with the F1 and R2 primers to generate the full length mutant construct SOE product. Mutant constructs were introduced into *V. cholerae* cells by chitin-induced natural transformation and/or cotransformation exactly as previously described (53, 54) or by chitin-independent transformations using a plasmid that ectopically expresses *tfoX* and *qstR* as previously described (55). For chitin-dependent natural transformation, *V. cholerae* was grown to mid-log in LB medium, washed with instant ocean medium (7 g/L; Aquarium Systems), and incubated with chitin flakes (Alfa Aesar) at final OD_600_ of 0.1 overnight at 30°C. The next day, SOE PCR products were added to chitin induced cells, incubated at 30°C for 5 hrs, and then outgrown and plated on selective media. For chitin-independent transformation, cells harboring pMMB67EH-*tfoX*-*qstR* were grown overnight in LB medium supplemented with<u>100</u> µM IPTG and 1 µg/mL chloramphenicol. The next day, 7 µL of the overnight culture was diluted into 350 µL of instant ocean medium. Then, SOE products were added as described above for chitin-dependent transformations. Mutant strains were confirmed by colony PCR, MAMA PCR (56), and/or sequencing. A complete list of strains and primers used in this study are outlined in **Table 1** and **Table 2**, respectively.

### Measuring GFP and mCherry fluorescence in bulk populations

GFP and mCherry fluorescence in reporter strains was determined exactly as previously described (57). Briefly, single colonies were grown in LB overnight at 30°C. Where indicated, media was supplemented with 10 µM CAI-1. CAI-1 was synthesized exactly as previously described (58). The next day, cells were washed and resuspended to an OD_600_ of 1.0 in instant ocean medium (7 g/L; Aquarium systems). Fluorescence was determined on a BioTek H1M plate reader. For GFP measurements, excitation was set to 500 nm and emission was set to 540 nm; for mCherry measurements, excitation was set to 580 nm and emission was set to 610 nm.

### Antibody generation

Purified *Vibrio harveyi* LuxR protein (300 µg; purified as previously described (59)) was sent to Cocalico Biologicals Inc. for serial injection into a rabbit host for antibody generation. Serum obtained from the third bleed has been used for Western analyses.

### Western blotting

From overnight cultures, cells were concentrated to an OD_600_ of 20 in instant ocean medium. Cells were lysed on a FastPrep-24 Classic Instrument at 4°C, then lysates were clarified by centrifugation. Lysates were then boiled with an equal volume of 2x SDS PAGE sample buffer (220 mM Tris pH 6.8, 25% glycerol, 1.2% SDS, 0.02% Bromophenol Blue, and 5% β-mercaptoethanol). Proteins were separated on a 15% SDS polyacrylamide gel by SDS electrophoresis, electropheretically transferred to a PVDF membrane, and probed with rabbit polyconal α-LuxR serum or mouse monoclonal α-RpoA (Biolegend) primary antibodies. LuxR is the HapR homolog in *Vibrio harveyi* and has 72% identity and 86% similarity to *V. cholerae* HapR. The LuxR antibody was empirically found to be cross-reactive with *V. cholerae* HapR, and so it was used to detect HapR protein levels. Blots were then incubated with α-rabbit or α-mouse HRP conjugated secondary antibodies, developed using Pierce ECL Western Blotting Substrate (ThermoFisher), and exposed to film.

### Chitin bead culturing

Chitin beads (200 µL of a 50% slurry; NEB) and overnight cultures of the indicated strains were washed with defined artificial salt water medium (DASW), which was prepared exactly as previously described (8). Chitin beads were inoculated with *V. cholerae* cells to an OD_600_ of 0.1 in a final volume of 5 mL in a Costar 6-well plate (Corning). Chitin mixtures were incubated statically at 30°C for 7 days before imaging.

### Microscopy data collection and analysis

To image chitin beads, cultured beads were gently transferred to a coverslip using wide-bore pipette tips. To image chitin grown cells, chitin bead reactions were vortexed then centrifuged at 250 x g for 1 minute. Cells found in the supernatant were transferred to a coverslip. To image Δ*cbp* strains, overnight cultures grown in LB were washed and resuspended to an OD_600_ of 0.2 in DASW and then transferred to a coverslip. Samples on the coverslip were placed under a 0.2% gelzan pad and imaged on an inverted Nikon Ti-2 microscope with a Plan Apo 60x objective, YFP & mCherry filter cubes, a Hamamatsu ORCAFlash 4.0 camera, and Nikon NIS Elements imaging software.

The strains used to examine P_*chb*_ expression across a population contained 1) a P_*chb*_-mCherry reporter and 2) a reporter that drove constitutive expression of GFP, P_*const2*_-GFP. By using these reporters, the expression of the P_*chb*_-mCherry could be normalized to GFP expression in each cell. Images were analyzed on Fiji using the MicrobeJ plugin (60) to determine mean intensity of cells in the YFP and mCherry channels. GFP was assessed using a YFP filter set to avoid background fluorescence from chitin beads, which was stronger in the GFP channel. Background fluorescence was subtracted from each channel and the mCherry / GFP fluorescence was determined for each individual cell.

### HapR protein purification

A plasmid expressing a hexahistidine-tagged *hapR* wild-type allele was generated using Gibson assembly. The *hapR* insert was amplified from *V. cholerae* E7946 and inserted into a pET28b vector (Novagen) using the primers listed in **Table 2**. The plasmid was then electroporated into *E. coli* BL21(DE3) for protein overexpression. This strain was grown overnight in LB medium with kanamycin, back-diluted 1:100 into 1 L of LB medium with kanamycin, and grown to an OD_600_ of 0.4-0.6 at 30°C. Expression of HapR was induced by IPTG to a final concentration of 1 mM and cultures grown for 4 h shaking at 30°C. The cells were pelleted and frozen at −80°C. The pellet was resuspended in 25 mL Buffer A (25 mM Tris pH 8, 500 mM NaCl), and an Avestin EmulfiFlex-C3 emulsifier was used to lyse cells. The soluble lysate was applied to a HisTrap HP Ni-NTA column using an Äkta Pure FPLC in Buffer A and eluted from the column with a gradient of Buffer B (25 mM Tris pH 8, 500 mM NaCl, 1 M imidazole). The purified protein was concentrated to approximately 5 ml using Sartorius Vivaspin Turbo 10,000 MWCO centrifugal concentrators. The sample was manually injected into the Äkta Pure and separated via size exclusion chromatography on a HiLoad™ 16/600 Superdex™ 75 pg column equilibrated with gel filtration buffer (25 mM Tris pH 7.5, 200 mM NaCl). Eluted fractions were analyzed by SDS-PAGE (15% gel), pooled, and concentrated using the same centrifugal concentrators previously mentioned. The samples were then immediately frozen in liquid nitrogen with a final concentration of 10% glycerol and stored at −80°C.

### Electrophoretic mobility shift assay (EMSA)

Binding reactions contained 10 mM Tris HCl pH 7.5, 1 mM EDTA, 10 mM KCl, 1 mM DTT, 50 μg/mL BSA, 0.1 mg/mL salmon sperm DNA, 5% glycerol, 1 nM of a Cy5 labeled DNA probe, and HapR at the indicated concentrations (diluted in 10 mM Tris pH 7.5, 10 mM KCl, 1 mM DTT, and 5% glycerol). Reactions were incubated at room temperature for 20 minutes in the dark, then electrophoretically separated on polyacrylamide gels in 0.5x Tris Borate EDTA (TBE) buffer. Gels were imaged for Cy5 fluorescence on a Typhoon-9210 instrument. Short DNA probes (30bp) were made by end-labeling one primer of a complementary pair (primers listed in **Table 2**) using 20 μM Cy5-dCTP and Terminal deoxynucleotidyl Transferase (TdT; Promega). Complementary primers (one labeled with Cy5 and the other unlabeled) were annealed by slow cooling at equimolar concentrations in annealing buffer (10 mM Tris pH 7.5 and 50 mM NaCl). P_*chb*_ and P_*clpP*_ probes were made by Phusion PCR, where Cy5-dCTP was included in the reaction at a level that would result in incorporation of 1–2 Cy5 labeled nucleotides in the final probe as previously described.

### ChIP-qPCR assays

Assays were carried out exactly as previously described (10). Briefly, overnight cultures were diluted to an OD_600_ of 0.08 and then grown for 6 hours at 30°C. Cultures were crosslinked using 1% paraformaldehyde, then quenched with a 1.2 molar excess of Tris. Cells were washed with PBS and stored at −80°C overnight. The next day, cells were resuspended in lysis buffer (1x FastBreak cell lysis reagent (Promega), 50 μg/mL lysozyme, 1% Triton X-100, 1 mM PMSF, and 1x protease inhibitor cocktail; 100x inhibitor cocktail contained the following: 0.07 mg/mL phosphoramidon (Santa Cruz), 0.006 mg/mL bestatin (MPbiomedicals/Fisher Scientific), 1.67 mg/mL AEBSF (DOT Scientific), 0.07 mg/mL pepstatin A (Gold Bio), 0.07 mg/mL E64 (Gold Bio)) and then lysed by sonication, resulting in a DNA shear size of ∼500 bp. Lysates were incubated with Anti-FLAG M2 Magnetic Beads (Sigma), washed to remove unbound proteins, and then bound protein-DNA complexes were eluted off with SDS. Samples were digested with Proteinase K, then crosslinks were reversed. DNA samples were cleaned up and used as template for quantitative PCR (qPCR) using iTaq Universal SYBR Green Supermix (Bio-Rad) and primers specific for the genes indicated (primers are listed in **Table 2**) on a Step-One qPCR system. Standard curves of genomic DNA were included in each experiment and were used to determine the abundance of each amplicon in the input (derived from the lysate prior to ChIP) and output (derived from the samples after ChIP). Primers to amplify *rpoB* served as a baseline control in this assay because HapR does not bind this locus. Data are reported as ‘Fold Enrichment’, which is defined as the ratio of the test promoter (P_*chb*_ or P_*clpP*_) / *rpoB* found in the output divided by the same ratio found in the input.

### Chitobiose competition

Overnight cultures grown in LB were washed with M9 medium and mixed 1:1. For each growth reaction, 10^2^ cells of this mixture was added to M9 medium supplemented with 0.2% chitobiose and 10 µM synthetic CAI-1 and grown shaking at 30°C for 24 hours. After 24 hours, ∼10^2^ cells from this mixture was used to inoculate fresh growth reactions to achieve additional generations of growth on chitobiose. This was repeated a third time after another 24 hours. CAI-1 was supplemented throughout these experiments to ensure consistently high levels of HapR expression throughout transfer steps. After each 24 hour growth period, the CFU/mL was determined for each strain in the mixture by dilution plating on LB agar supplemented with X-gal. Competing strains were discerned by blue/white screening as one strain was *lacZ*^+^ and the other was Δ*lacZ*. A *lacZ*^+^ strain was competed against a Δ*lacZ* strain (parent : parent competition) or against a Δ*lacZ* Δ*hapR* strain (Δ*hapR* : parent competition). *V. cholerae* can grow on chitobiose through the activity of two sugar transporters: the GlcNAc PTS transporter (VC0995) or the chitobiose ABC transporter (VC0616-0620).

Because we wanted to study the regulation of the transporter encoded within the *chb* locus, all strains for this assay had a deletion in VC0995, which renders growth on chitobiose dependent on the *chb* locus as previously described (11). Competitive indices were calculated as the CFU ratio of Δ*lacZ* / *lacZ*^+^ after growth for the indicated number of generations divided by the CFU ratio of the Δ*lacZ* / *lacZ*+ in the initial inoculum.

### Statistics

Statistical comparisons were determined using Student’s t-test or One-way ANOVA with Tukey’s post-test using GraphPad Prism software.

## Acknowledgements

We thank Ryan Chaparian for offering experimental advice, David Grainger for helpful discussions, and Wai-Leung Ng for providing synthetic CAI-1. This work was supported by grant R35GM128674 from the National Institutes of Health to ABD and R35GM124698 from the National Institutes of Health to JVK.

## Supplemental Information for

**Figure S1.**
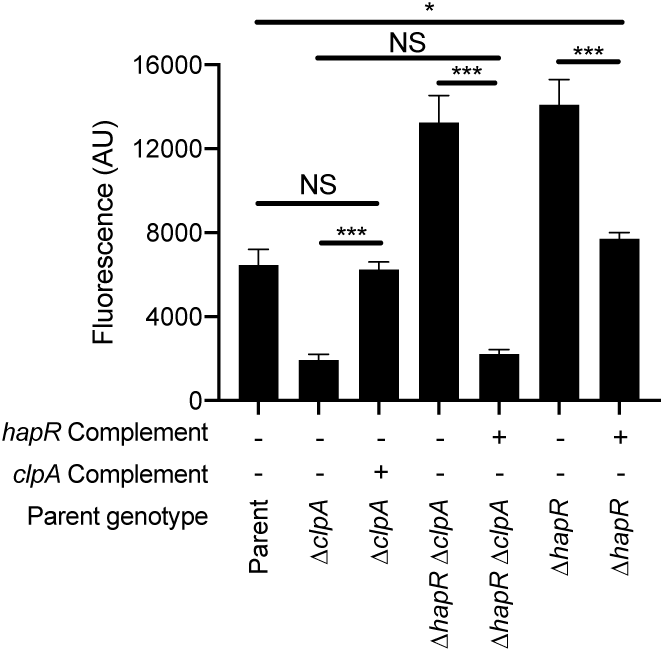
Complementation of ΔhapR and ΔclpA mutant strains. Expression of a P_*chb*_-GFP transcriptional reporter was determined in the indicated strains. The parent strain contained a chromosomally-encoded P_*chb*_-GFP reporter and a Δ*cbp* mutation. Strains were complemented with a chromosomally-encoded single copy of the indicated gene driven by its native promoter at an ectopic site (*hapR* or *clpA* Complement). Fluorescence of cultures was determined on a plate reader from at least three independent biological replicates and is shown as the mean ± SD. Statistical comparisons were made by one-way ANOVA with Tukey’s post-test. NS, not significant. *, p = 0.0104. ***, p < 0.001. The P_*chb*_-GFP fluorescence data for “Δ*clpA*”, “Δ*hapR* Δ*clpA*”, and “Δ*hapR*” are identical to the data presented in **Fig. 1** and were included here for ease of comparison.

**Figure S2.**
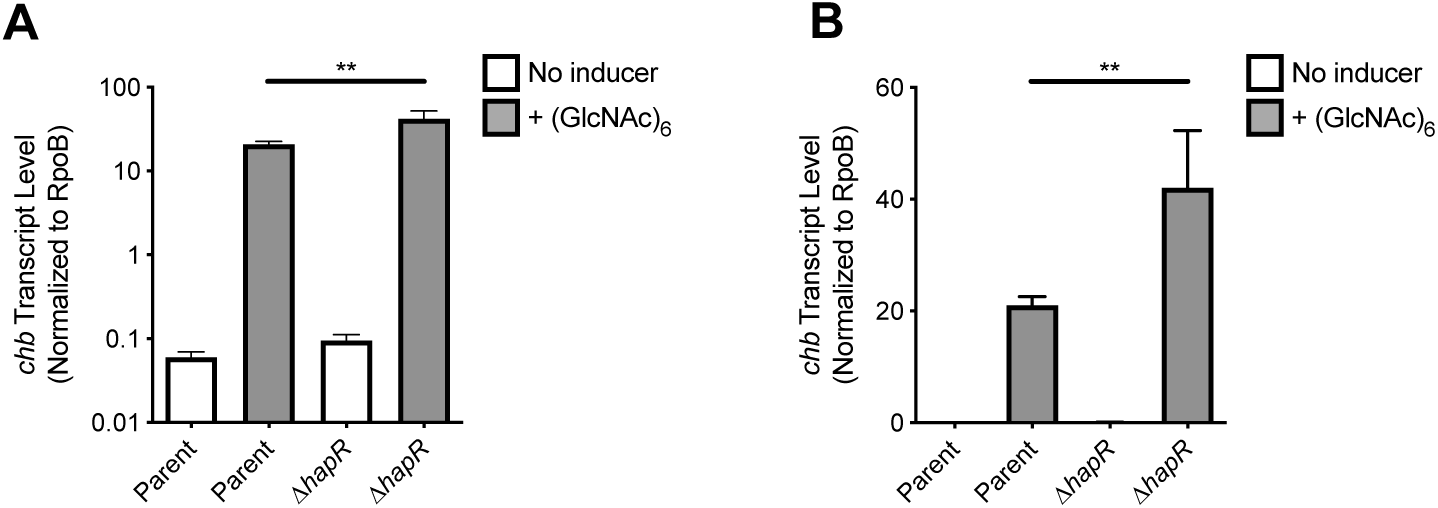
HapR is a repressor of native chb expression. The indicated strains were induced with sterile water (no inducer) or chitin oligosaccharides (+ (GlcNAc)_6_), then assessed for *chb* transcript abundance by qRT PCR. The parent strain used was *V. cholerae* E7946 WT. Data are shown on **A**) a log scale or **B**) a linear scale. The latter plot helps accentuate that deletion of *hapR* results in a two-fold increase in native *chb* transcripts in response to chitin oligosaccharides, which is similar to what is observed using transcriptional reporters. Data are from at least three independent biological replicates and is shown as the mean ± SD. Statistical comparisons were made by one-way ANOVA with Tukey’s post-test. **, p = 0.0064.

**Figure S3.**
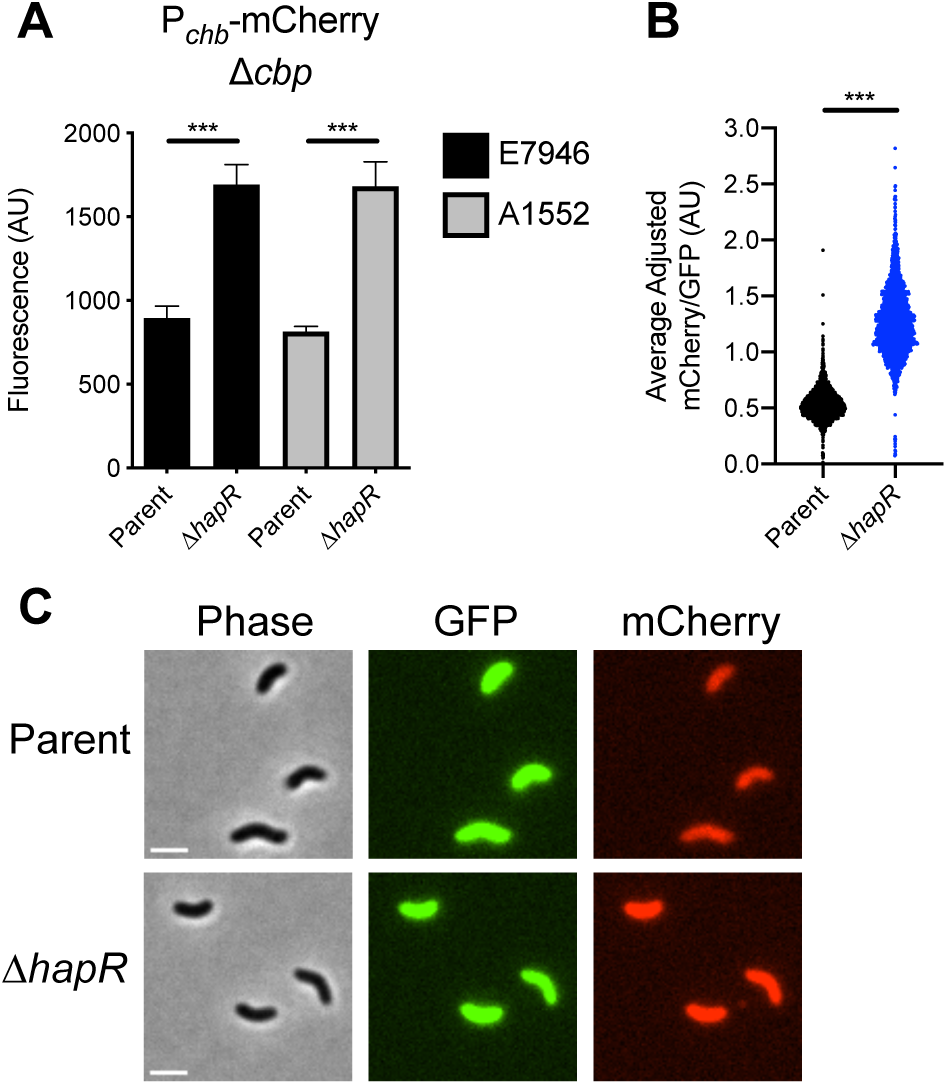
HapR is a repressor of P_chb_ in V. cholerae E7946 and V. cholerae A1552. **A**) Expression of a P_*chb*_-mCherry reporter was determined in the indicated mutant strains. Each respective parent strain contained a P_*chb*_-mCherry reporter and a Δ*cbp* mutation to activate ChiS. **B**) Scatter plot showing the relative expression of a P_*chb*_-mCherry reporter in the indicated *V. cholerae* A1552 strains that have *cbp* intact when using chitin oligosaccharides to induce cells. The parent strain background contains both a P_*chb*_-mCherry reporter, a P_const2_-GFP construct (which exhibits constitutive GFP expression), and *cbp* intact. n = 2189 for Parent; n = 2164 for Δ*hapR.* Data shown are from two independent biological replicates. **C**) Representative images of *V. cholerae* A1552 strains analyzed in **B**. Scale bar = 2 μm. Statistical comparisons in **A** were made using a one way ANOVA with Tukey’s post test and in **B** were made using Student’s t-test. ***, p < 0.001.

**Figure S4.**
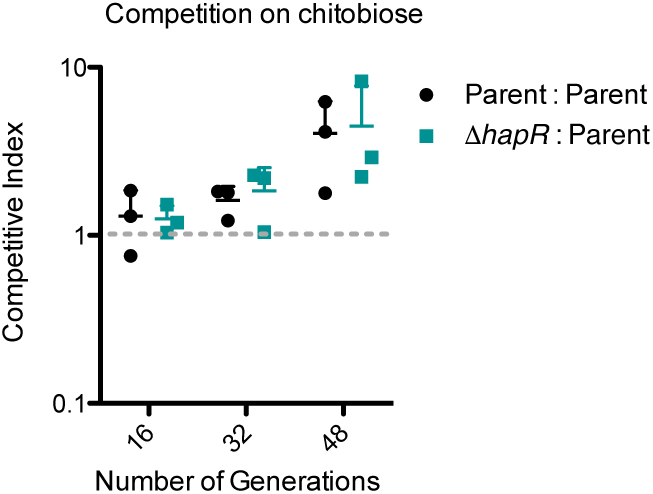
A ΔhapR mutant does not have a fitness advantage for growth on chitobiose. The indicated *V. cholerae* strains were mixed and co-cultured in M9 minimal media supplemented with 0.2% chitobiose + 10 µM CAI-1 and grown for the indicated number of generations. Each data point represents an independent biological replicate.

**Figure S5.**
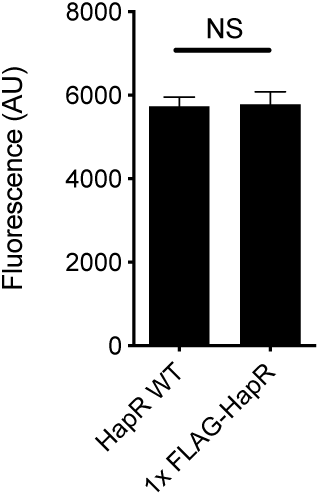
N-terminally FLAG-tagged HapR is functional for regulation of P_chb_. Strains expressing the indicated HapR allele at the native locus were assessed for expression of a P_*chb*_-GFP reporter. Both strains also harbored a Δ*cbp* deletion to activate ChiS. Statistical comparisons were made using Student’s t-test. NS, not significant.

**Figure S6.**
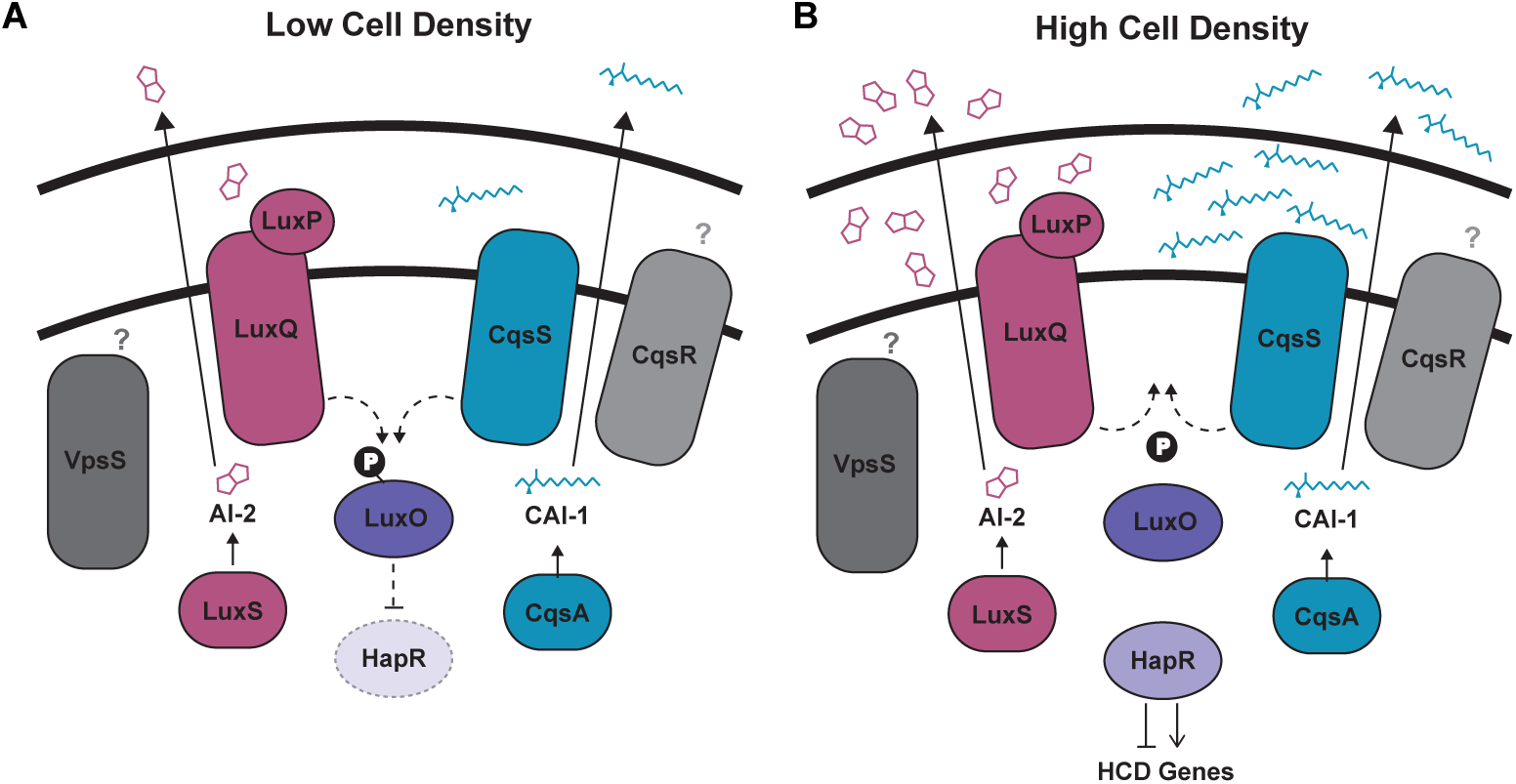
Model of quorum sensing regulation in *V. cholerae*. There are four parallel sensor kinases that contribute to QS in *V. cholerae*: LuxQ, CqsS, VpsS, and CqsR. The signals for VpsS and CqsR are unknown. LuxPQ senses the interspecies autoinducer AI-2 and CqsS senses the *V. cholerae*-specific autoinducer CAI-1. AI-2 is produced by the LuxS synthase and CAI-1 is produced by the CqsA synthase. (**A**) At low cell density, the LuxQ and CqsS sensors act as kinases, resulting in phosphorylation of LuxO, which subsequently decreases expression of HapR. (**B**) At high cell density, LuxQ and CqsS act as phosphatases, which results in dephosphorylation of LuxO, and subsequent activation of HapR expression. Dashed lines indicate indirect effects of one protein on another.

**Table S1.**
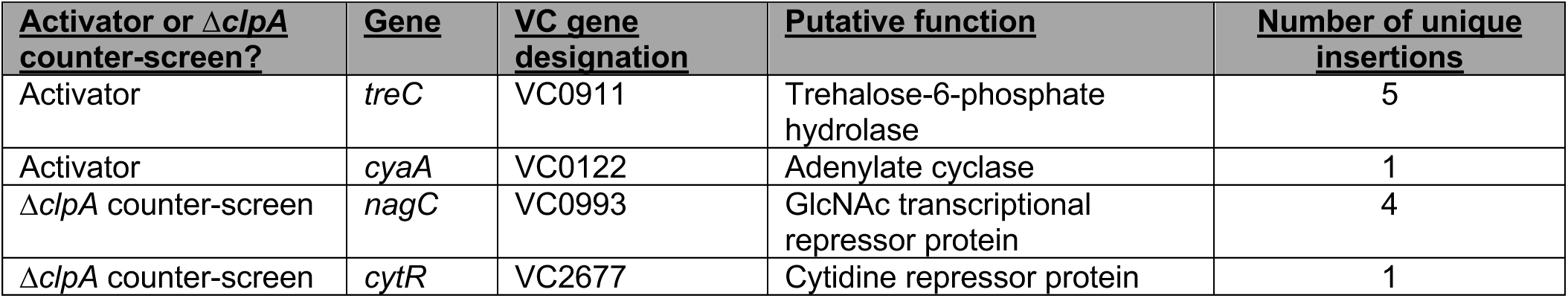
Other hits from P_chb_ activator screen and ΔclpA counter-screen.

